# Competition NMR for Detection of Hit/Lead Inhibitors of Protein-Protein Interactions

**DOI:** 10.1101/2020.06.12.148379

**Authors:** Bogdan Musielak, Weronika Janczyk, Ismael Rodriguez, Jacek Plewka, Dominik Sala, Katarzyna Magiera-Mularz, Tad A. Holak

**Affiliations:** Department of Organic Chemistry, Faculty of Chemistry, Jagiellonian University, Gronostajowa 2, 30-387, Krakow, Poland; Max Planck Institute for Biochemistry, Am Klopferspitz 18, 82152 Martinsried, Germany

## Abstract

Screening for small-molecule fragments that can lead to potent inhibitors of protein-protein interactions (PPIs) is often a laborious step as the fragments cannot dissociate the targeted PPI due to their low μM-mM affinities. Here, we describe an NMR competition assay - called w-AIDA-NMR (weak-Antagonist Induced Dissociation Assay-NMR) - that is sensitive to weak μM-mM ligand-protein interactions and which can be used in initial fragment screening campaigns. By introducing point mutations in the complex’s protein that is not targeted by the inhibitor, we lower the effective affinity of the complex allowing for short fragments to dissociate the complex. We illustrate the method with the compounds that block the Mdm2/X-p53 and PD-1/PD-L1 oncogenic interactions. Targeting the PD-/PD-L1 PPI has profoundly advanced the treatment of different types of cancers.

## INTRODUCTION

NMR spectroscopy is a highly versatile screening method for drug discovery^1–3,^. In comparison to other screening technologies, NMR is capable of detecting the binding of small-molecule compounds to macromolecular targets over an extraordinary broad affinity range from covalent to millimolar^4,5^. A unique feature of NMR is its robust capability to detect weak intermolecular interactions. This feature makes NMR ideal for fragment-based screening, in which the binary binding between low-affinity fragments and target proteins is studied^1–3,6–8^. This type of NMR “binary screening” does not provide information whether a compound can inhibit or dissociate protein-protein interactions. We have recently described an NMR-based assay for studying the effect of antagonists on protein-protein interactions^9–13^. The method, named AIDA-NMR (for the Antagonist Induced Dissociation Assay-NMR), belongs to the target protein-detected NMR screening methods^14^ and provides unambiguous information on whether an antagonist of a protein-protein interaction is strong enough to dissociate the complex. Whether a hit/lead compound is capable of dissociating a particular protein-protein interaction (PPI) is determined by the strength of its binding to one of the two protein components of the PPI. If the binding is comparable or stronger than the affinity of the protein-protein interaction, which commonly is at the low μM to nM range, the PPI complex should be broken up. The “weak” small-molecule binders to the proteins of the PPI; for example, those ligands with the IC_50_ in high μM and mM range, would not affect the AIDA-NMR (although there is, albeit weak, binding to one of the protein of the PPI). Thus, the strength of the PPI complex determines the weakest binding compounds that may still be measured with the AIDA-NMR.

Herein, we describe an AIDA-NMR technology that should overcome these disadvantages, allowing for NMR to be used in the primary fragment screening. We use the Mdm2-p53 and MdmX-p53 interactions^15,16^ as proof-of-concept systems to develop our protocol and use the method on the immune oncology system of the PD-1/PD-L1 interaction^17,18^. The tumor suppressor p53 protein is a key player in protecting the organism from cancer and was therefore coined a term: “the guardian of the genome. To escape the control system mediated by p53, majority of human cancers has either mutation within p53 (50% all cancers), whereas the rest compromises the effectiveness of the p53 pathway^19,20^. In tumors with unmodified wildtype p53, the p53 pathway is inactivated by the Mdm2 and MdmX proteins^21,22^. Therefore, the disruption of the Mdm2-p53/MdmX-p53 interactions that leads to the restoration of the impaired function of p53, poses a new approach to anticancer therapies across a broad spectrum of cancers.

Another important protein target in cancer is the PD-1/PD-L1 system. PD-1 (Programmed cell death protein 1) is expressed on activated T cells and plays a critical role in modulation of the host’s immune response^23,24^. The principal PD-1 ligand, PD-L1, is expressed on macrophages, monocytes and cancer cells. Cancer cells exploit this ligand protein to avoid immune attack by T cells^25^. This seminal finding of how cancer cells use binding between PD-L1 and PD-1 to inhibit the killing of tumor cells by T cell have now been translated into effective medical treatment^17,26–30^. Blocking the immune checkpoint PD-L1 or PD-1 allows for the T cell killing of tumor cells, and immune checkpoint inhibitors targeting the PD-1/PD-L1 interaction have revolutionized modern cancer therapy for advanced cancer^18,31–33^.

The interaction of PD-L1/PD-1 and p53-Mdm2/X present challenging cases in detecting weak binders, because the K_D_ of this interaction is 8.2 and 0.2-0.6 μM, respectively^34,35^. We show herein that by introducing designed mutations in the component protein(s) of these protein-protein complexes, we could weaken the affinity of the PPI interaction. The mutations are introduced within the binding partner that is not targeted with inhibitors, therefore not compromising the binding interface of targeted protein and tested ligand. For example, in the case of p53/Mdm2 complex, we have mutated p53 to determine fragments binding to Mdm2 that dissociates the complex. Since there are over 30 000 known naturally occurring missense mutations to p53^36,37^ such a system is physiologically still relevant. Mutations within the non-targeted binding partner allow for the proteins that build up the PPI complex to be sensitive to weak binding compounds i.e. the modified assay can be used in the fragment-based screening. We named this variant of the AIDA experiment “w-AIDA-NMR”, where “w” stands for “weak”.

## RESULTS

### p53 mutants

The complexes of wt-p53 with Mdm2 and MdmX have K_D_ values 0.60 μM and 0.24 μM, respectively^35^. These K_D_’s determine the weakest inhibitor, which can be tested with the AIDA-NMR methods. For the compounds that weakly bind to Mdm2/X, which still could be good initial scaffolds for further optimization in drug development process, the method would not be sensitive enough for detecting inhibition of the p53-Mdm2 and p53-MdmX PPI.

Three key residues Phe19, Trp23 and Leu26 of p53 make the highest contribution to the binding energy of p53 with Mdm2/X^38–40^. Among them, Trp23 is the most important, and any mutations of this residue completely abrogate the binding between p53 and Mdm2/X^39^. Mutations of Phe19 have similar effects, but not as strong as those of Trp23. Both residues are essential for the p53/Mdm2 complex formation. They are buried within pockets of Mdm2/X with strong π-π stabilizing interactions (Fig. 1a,b; Supplementary Fig. 1)^39^. Since the aim of the research was not to block the interaction, but only to slightly weaken it, Phe19 and Trp23 were not touched, and Leu22 and Leu26 were chosen as most plausible targets of the mutations (Fig. 1a,b; Supplementary Fig. 1). In both Mdm2 and MdmX complexes with p53, the side chains of Leu22 and Leu26 are pointing to the outside of the main pocket occupied by Trp23 and Phe19 and had little effect on the structural arrangements of the p53-binding pockets of Mdm2/X, making them ideal candidates for our w-AIDA assay^38,40,41^ (Fig. 1a, b and Supplementary Fig. 1).

**Fig. 1.**
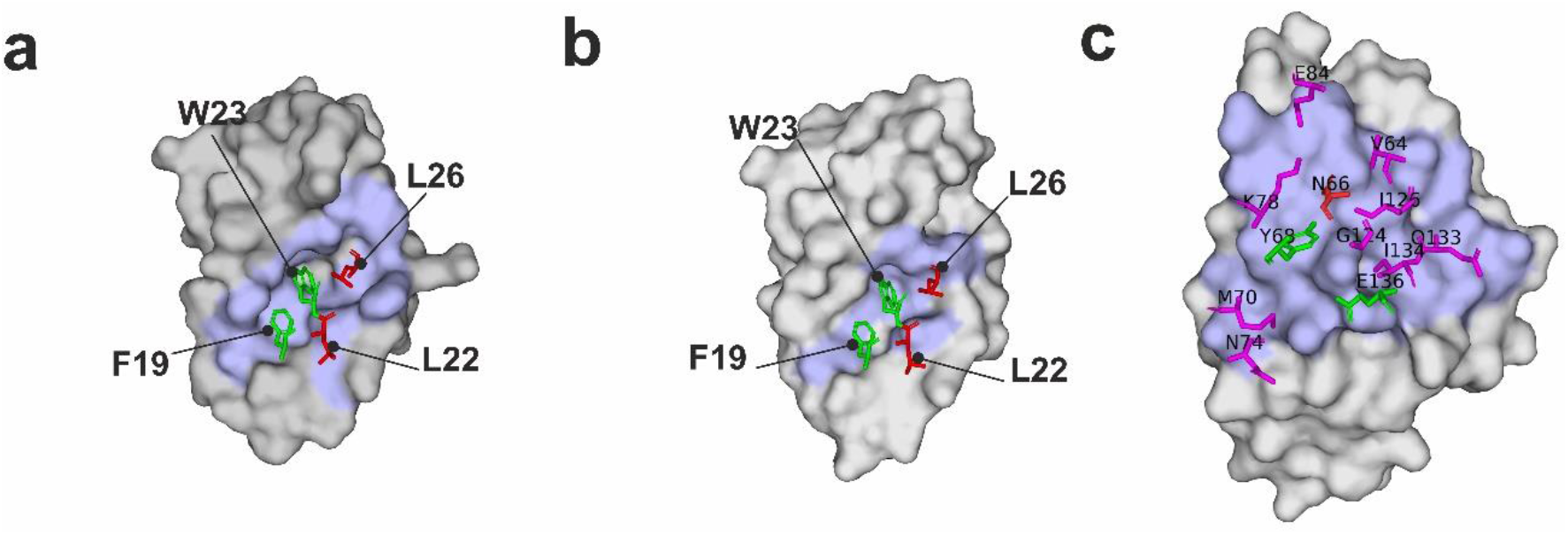
Visualization of the p53/Mdm2 (a), p53/MdmX (b) and PD-1/PD-L1 (c) complexes (PDB IDs:1YCR, 3DAB and 4ZQK, respectively). The surface representation of Mdm2, MdmX and PD-L1 is shown in gray with their interfaces with p53 and PD-1 in pale blue. For better visibility of the interfaces, p53 peptide is fully depicted as a cartoon, while only crucial for interactions PD-1 residues are shown. Interacting residues from p53 and PD-1 are shown as sticks with purple one interacting with binding partner, but not being essential for the interactions, green being residues whose mutation abolishes the interactions and red being amino acids that were mutated within this manuscript consideration.

Taken together, these small changes in the p53 sequence should result in shortening of the side chain (L→A, L→V) or changing of the amino acids isoform (L→I). K_D_ values for the complexes of Mdm2/X with the p53 mutants were established with the isothermal titration calorimetry (ITC) (Table 1, Supplementary Figs. 2-8).

**Table 1.**
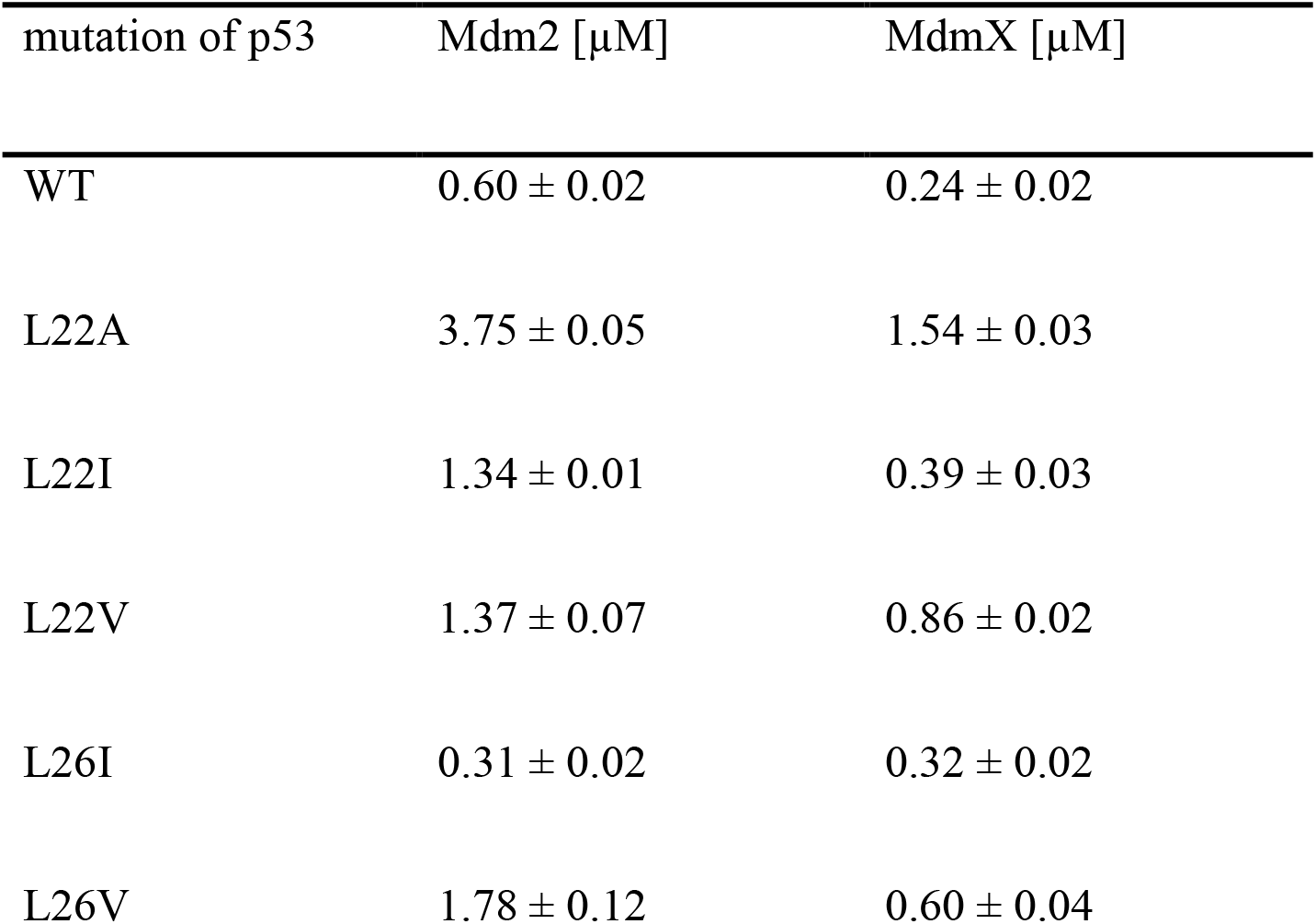
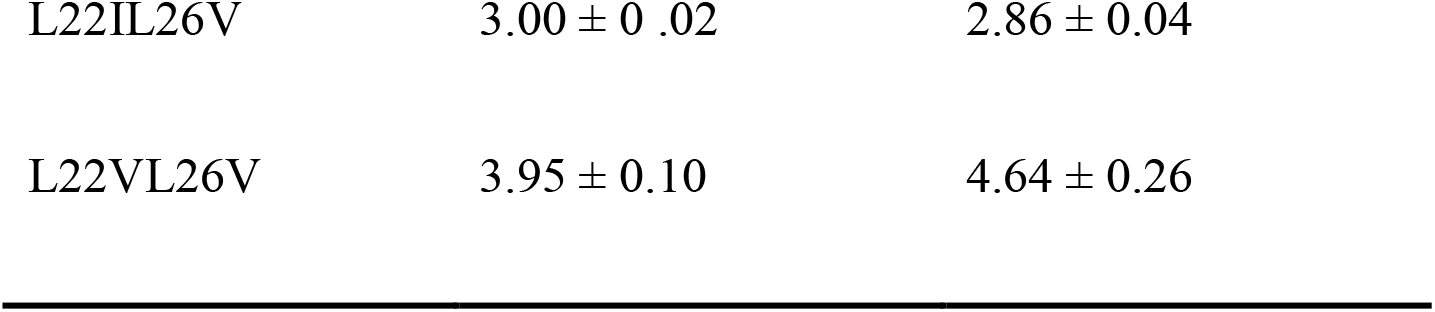
Dissociation equilibrium constants of p53-wt and p53 mutants with Mdm2 and MdmX determined using ITC. Fitted curves are shown in Supplementary Figure 2-8.

| mutation of p53 | Mdm2 [μM] | MdmX [μM] |
| --- | --- | --- |
| WT | $0.60 \pm 0.02$ | $0.24 \pm 0.02$ |
| L22A | $3.75 \pm 0.05$ | $1.54 \pm 0.03$ |
| L22I | $1.34 \pm 0.01$ | $0.39 \pm 0.03$ |
| L22V | $1.37 \pm 0.07$ | $0.86 \pm 0.02$ |
| L26I | $0.31 \pm 0.02$ | $0.32 \pm 0.02$ |
| L26V | $1.78 \pm 0.12$ | $0.60 \pm 0.04$ |
| L22IL26V | $3.00 \pm 0.02$ | $2.86 \pm 0.04$ |
| L22VL26V | $3.95 \pm 0.10$ | $4.64 \pm 0.26$ |

The most significant lowering of the Mdm2/X-p53 interaction was observed for the L22A mutation, where the K_D_ value increased six fold compared to that of the p53-wt/Mdm2 complex. Surprisingly, modifications of Leu26, which is one of the key amino acids of the p53 binding to Mdm2/X, were less efficient than mutations of Leu22. Moreover the L26I mutant generated lower K_D_ value (0.31 μM) than in the Mdm2/p53-wt interaction (0.60 μM). Although the interaction with the L26V mutant was weakened, the affinities of the proteins were still too strong to use the preformed complexes for the investigation of the weak binding compounds. Therefore, we designed the double mutants (L22IL26V and L22VL26V) which combine above-mentioned mutations. The ITC measurements showed a synergistic effect of the double mutations on the binding of the mutant p53s to Mdm2/X (Supplementary Figs. 7 and 8). One-dimensional proton NMR spectra of all mutants were almost identical to that of the wt-p53 spectra. This indicated that the mutated p53 constructs were correctly folded (data not shown).

### Inhibitors of the wt-p53/Mdm2 and wt-p53/MdmX interactions

A large number of low-molecular-weight compounds that bind to Mdm2 and MdmX^42–44^ have been tested in clinical trials for several years^45,46^. Among the most advanced are cis-imidazoline derivatives called nutlins^47^. **Nutlin-3a** is a selective and potent inhibitor of the p53-Mdm2 interaction. The nutlin-3a derivative idasanutlin, is currently under investigation in trials involving patients with advanced-stage haematological malignancies or solid tumors^48^.

Three representative and well-studied small-molecule inhibitors of the wt-p53/Mdm2 interaction^49^ (Fig. 2) and three for the wt-p53/MdmX^50^ (Fig. 3) were chosen to test their inhibition of the complexes: the wt-p53/Mdm2, mutant-p53/Mdm2, wt-p53/MdmX, and mutant-p53/MdmX.

**Fig. 2.**
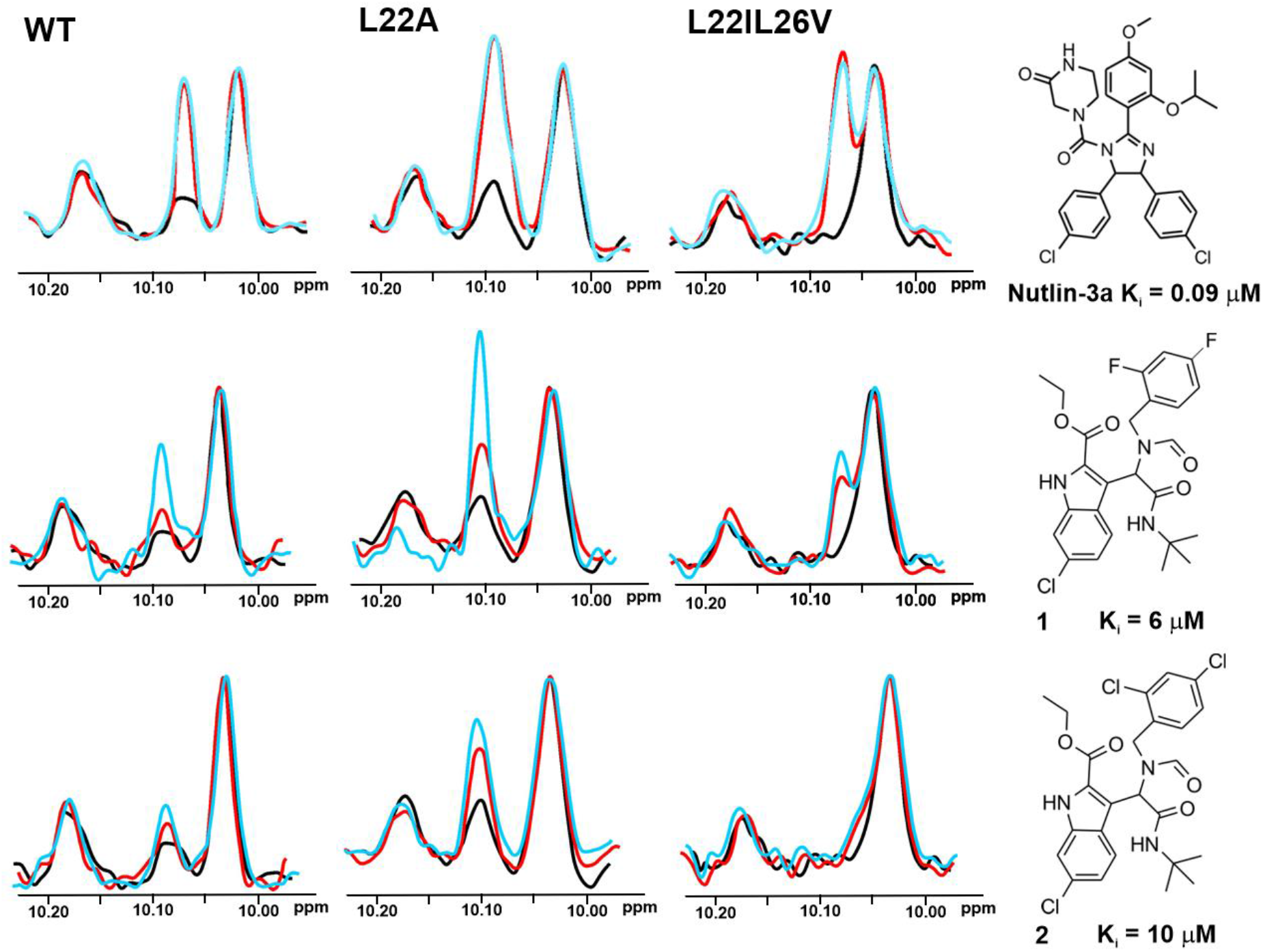
The 1D AIDA proton NMR spectra of the side chain ^N^H^∊^ protons of tryptophans for the p53/Mdm2 complexes (p53-wt, L22A and L22IL26V). All protein complexes (black) were treated with **Nutlin-3a**, compounds **1** and **2** in molar ratio: the protein to a compound 1:1 (red) and 1:2 (blue), respectively.

**Fig. 3.**
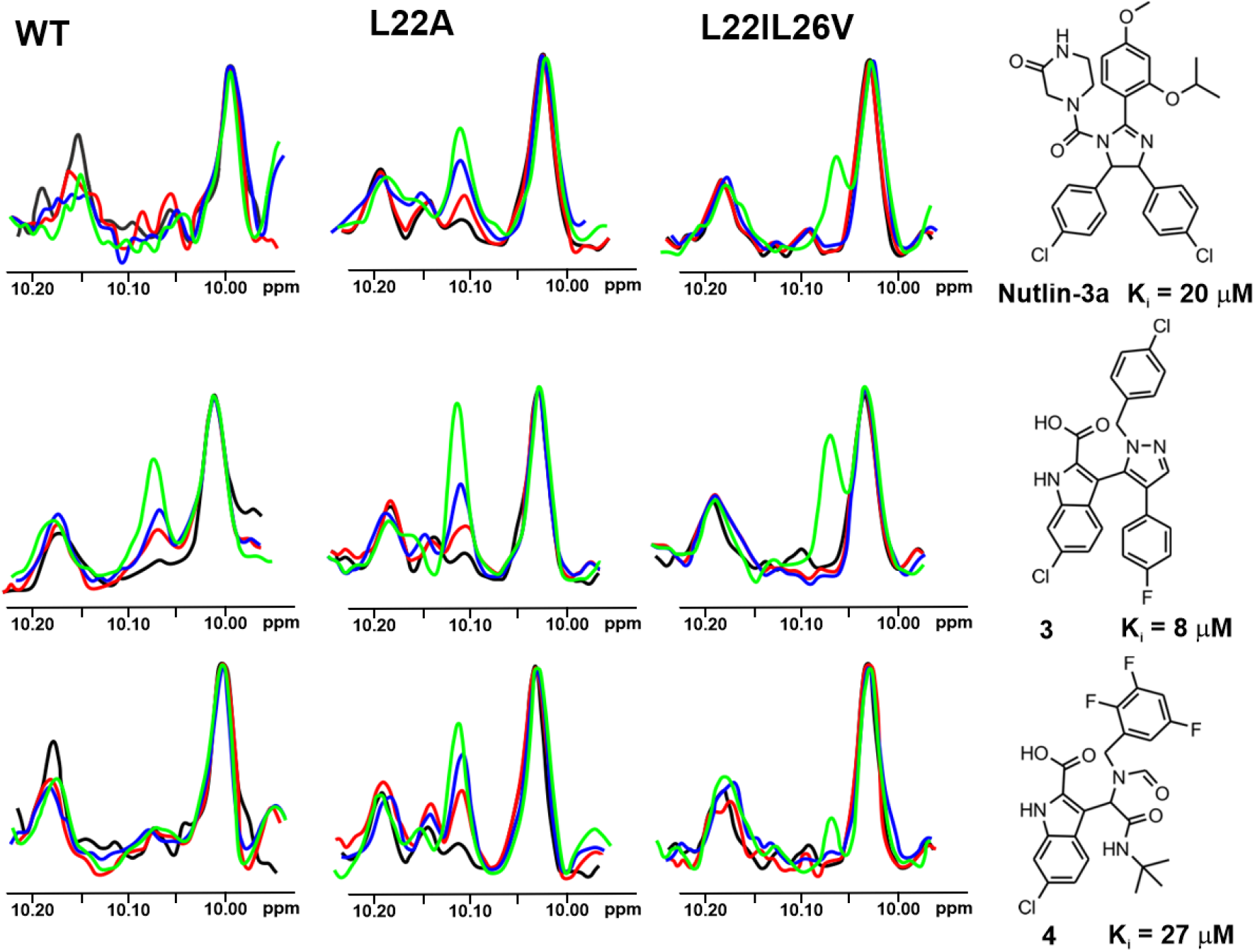
The 1D AIDA proton NMR spectra of the side chain ^N^H^∊^ proton of tryptophans for the mutant-p53/MdmX complexes (p53-wt, L22A and L22IL26V). All protein complexes (black) were treated with **Nutlin-3a**, compound **3** and **4** in molar ratio: the protein to a compound 1:1 (red), 1:2 (blue) and 1:5 (green), respectively.

### AIDA-NMR for the wt-p53/Mdm2 and wt-p53/MdmX complexes

The AIDA-NMR assay, in its 1D ^1^H NMR version^13,14,51^, was carried out with all selected compounds on the wt-p53 complexes. The 1D AIDA NMR experiment is based on monitoring the recovery of the signal of the ^N^H^ε^ proton of Trp23 upon addition of an inhibitor (Supplementary Fig. 9)^13,14,51^. The full recovery of this signal was not observed for these five inhibitors even with the five-fold molar excess of the compound over the proteins complex. Only two inhibitors **1** (K_i_ value 6 μM towards Mdm2) and **3** (K_i_ value 8 μM towards MdmX) showed some weak interactions (Supplementary Fig. 10). That means that the K_i_ values of tested compounds are too high and the method that uses wt-p53 is not sensitive enough for monitoring these weaker binders to Mdm2/X.

### w-AIDA-NMR for the p53/Mdm2 and p53/MdmX complexes

To choose proper mutants for the 1D NMR w-AIDA assay, we recorded ^1^H NMR spectra of the all designed mutants (Fig. 4). Since the 1D AIDA NMR experiment is based on the observation of the recovery of the signal of Trp23 upon addition of an inhibitor (Supplementary Fig. 9)^13,14,51^, it was crucial that the change in the position of that signal does not lead to it being obscured by overlapping with other signals. Although the introduced mutations were not significant for the overall protein structure, small changes in the local magnetic environment of Trp23 could be observed. These changes resulted in a slight shift of a position of the NMR signal corresponding to the ^N^H^ε^ proton of Trp23 (L22A, L22I), partial overlapping (L22V, L26V, L22IL26V) or even the complete overlapping of the Trp23 with Trp53 signals (L22VL26V).

**Fig. 4.**
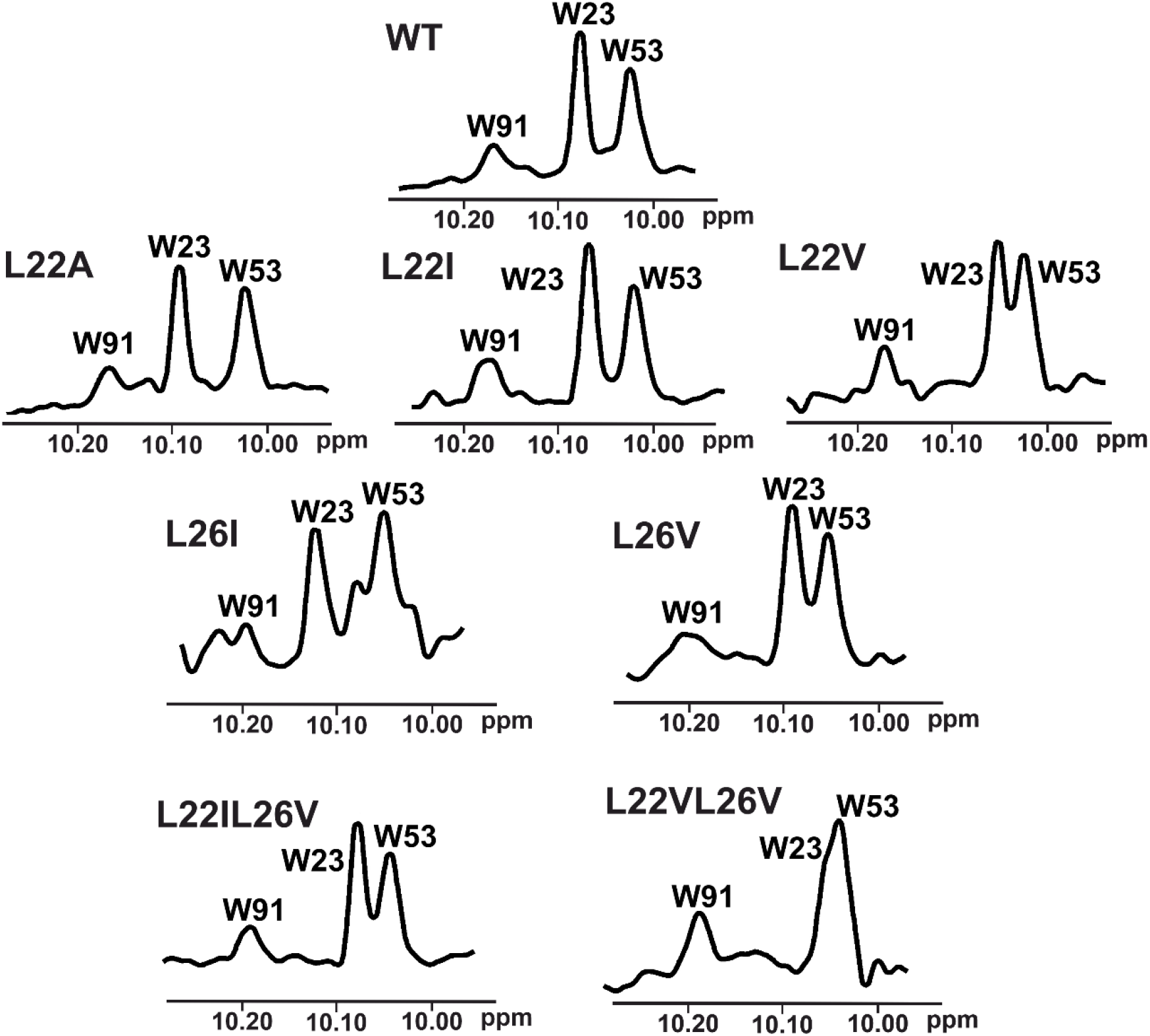
An enlarged part of the ^1^H NMR spectrum of wt-p53 and its mutants. Visible peaks originate from the side chain ^N^H^∊^ protons of W91, W23, and W53 left to right, respectively.

Based on the affinities determined with the ITC experiments (Supplementary Table 1 and Supplementary Figs. 2-8), as well as ^1^H NMR spectra of the mutants (Fig. 4), the L22IL26V and L22A mutants were chosen for evaluating their application in the ^1^H w-AIDA-NMR assay. Application of mutants L22A and L22IL26V increased the sensitivity of 1D AIDA-NMR assay. The Mdm2 inhibitors **1** and **2** caused almost full recovery of the Trp23 signal with the L22A mutant (Fig. 2). For MdmX, all tested MdmX inhibitors released the p53 from its complex with MdmX (Fig. 3). Better results were obtained for the L22A mutant than for the L22II26V mutant.

### PD-1 mutants

The second PPI that we tested with the w-AIDA-NMR is the interaction of human PD-L1 with PD-1. Here we use the entire ectodomain of PD-1 and the PD-1 binding domain of the ectodomain of PD-L1 (residues 18-134) (Supplementary Fig. 11). Although the PD-1/PD-L1 complex is characterized by a relatively high affinity constant (Supplementary Fig. 12), it is, nevertheless, too strong to be used in the NMR screening of “weak” fragments, which usually have two or three orders of magnitude lower affinities. Therefore, we designed a series of the PD-1 mutants based on the structure of the PD-1/PD-L1 complex^52^. We identified the amino acid candidates on PD-1 for the mutations using the same approach as that described for the p53/Mdm2/MdmX systems. The mutations should weaken the binding of PD-1 to PD-L1 in a minimal invasive manner (Fig. 1c, Supplementary Figs. 1 and 13). The mutated PD-1 N66A, Y68A, E135A and the double mutant N66AY68A were then expressed in *E. coli* (Supplementary Table 1) and checked by NMR whether the mutations affected the folding of the protein. From these tested mutants only the N66A mutant was folded (the data for other mutants not shown). Protein melting analysis showed that the N66A PD-1 mutant has similar midpoint of thermal transition (melting temperature), indicating that the changes in its stability against denaturation were insignificant; in fact, the N66A mutant was slightly more temperature stable than wt-PD-1 (Supplementary Fig. 14). The affinities of the PD-1/PD-L1 and (N66A)PD-1/PD-L1 complexes was determined using the Microscale Thermophoresis by titrating the labeled PD-1 with the unlabeled PD-L1. The resulting data was next fitted with K_D_ models to yield affinities. For the (N66A)PD-1/PD-L1 complex, we were not able to reach the top plateau as we could not concentrate PD-L1 solution above 183 μM (Supplementary Fig. 12). Therefore, the K_D_ is estimated to be above 100 μM, considerably higher than for the native complex, allowing weaker small-molecular fragments to dissociate it. The N66A mutation did not affect the overall structure of the interface of the PD-1/PD-L1 complex as depicted in the Supplementary Fig. 13. Asn66 is only involved in one hydrogen bond with Ala121 from PD-L1 and when mutated to alanine does not significantly change the space in the complex (Supplementary Fig. 13 inset). Moreover, when overlaid on top of each other the critical for interactions residues from hPD-L1 are virtually in the same position regardless to the mutations of PD-1 including murine PD-1-human PD-L1 complex (Supplementary Fig. 13).

### Inhibitors of the immunocheckpoint PD-1/PD-L1 interaction

A sizeable number of small-molecule binders to PD-L1 that inhibit this PPI has now been described (ca. 1000)^53,54^. We have recently published a series of studies on the affinities of the small-molecule inhibitors of the PD-1/PD-L1 interaction developed by Bristol-Myers Squibb (called herein the BMS compounds^55,56^). The BMS compounds are based on the hydrophobic biphenyl core scaffold^53,57^. Our NMR studies indicate that the binding of the BMS compounds to PD-L1 induces oligomerization of the protein^57–59^. The line width broadening in the NMR signals of PD-L1 after addition of BMS compounds, both in proton and ^1^H-^15^N 2D HMQCs is so extensive that it is impossible to estimate dissociation constants also for “weaker” binding precursors of the BMS compounds (K_D_’s double digit μM). One of the NMR techniques that would allow validating the binding and estimating the value of K_D_ is the w-AIDA-NMR assay.

For the start in the PD-1/PD-L1 system, we used two smallest fragments of the **BMS-1166** inhibitor (fragments **5** and **6** in Supplementary Fig. 15)^59^. **BMS-1166** is one of the most potent small-molecule ICB inhibitors for the PD-1/PD-L1 developed until now^53^. It binds to PD-L1 and efficiently dissociates the human PD-1/PD-L1 complex *in vitro*^59^. In the ICB cell models, it activates the effector T cells, which are attenuated by both soluble and membrane-bound PD-L1 presented by antigen-presenting cells^59^.

### AIDA-NMR on the wt-PD-1/wt-PD-L1 complex

For the start in the PD-1/PD-L1 system, we used two smallest fragments of the **BMS-1166** inhibitor (fragments **5** and **6** in Supplementary Fig. 15) that in our previous tests showed the interaction with the PD-L1 protein in the ^1^H-^15^N HMQC NMR^59^. To perform the 2D AIDA experiment for **5** and **6**, as the so-called reporter protein, which should be ^15^N isotopically labeled, we use the ^15^N-labeled PD-1 (13.2 kDa) (Supplementary Fig. 16a). After addition of PD-L1 (14.9 kDa) in the molar ratio 1:1, most of the cross-peaks in the ^1^H-^15^N HMQC spectrum of PD-1 became broader, their intensities decreased and most of the cross peaks disappeared (Supplementary Fig. 16b). This result confirms the forming of the complex with the molecular weight ca. 30 kDa. The AIDA-NMR assay was then applied to test the dissociating capabilities of **5** and **6**. The compounds that despite being “active” in the binary interaction with PD-L1, were “inactive” in the AIDA test. No recovery signals from of the ^15^N labeled PD-1 protein was observed in 2D and 1D spectra (Supplementary Figs. 16c-d and 17). This shows that the tested compounds are not able to dissociate the PD-1/PD-L1 complex; their dissociation constants with PD-L1 are higher than dissociation constant of the native human PD-1/PD-L1 complex (8 μM – Supplementary Fig. 12). We also performed a positive control and the full recovery of 2D HMQC spectrum of the ^15^N PD-1 was observed after adding **BMS-1166** with K_D_ = 1.4 nM to sample with compound **5** (Supplementary Fig. 16e).

### w-AIDA-NMR for the (N66A)PD-1/PD-L1 complex

In the same way as for the wt-PD-1 proteins, we performed an 2D AIDA-NMR experiment using the ^15^N-labeled mutated protein (N66A)PD-1 (13.2 kDa) and wt-PD-L1 (Figs. 5a and 5b). In contrast to the pervious experiment, the resulting complex of the (N66A)PD-1/PD-L1 proteins (28.1 kDa) has smaller number of broadening/disappearing signals in the HMQC spectra due to the higher K_D_ value. However, most noticeable changes of the chemical shifts can be observed in the range ca. 8.8-9.4 and 122-127 ppm for hydrogen and nitrogen, respectively (Fig. 5b). Addition of an equimolar amount of **5** or **6** results in dissociating of the PD-1/PD-L1 complex (Figs. 5c and 5d) with appreciable recovery of the 2D signals observed for **5**. This suggests that the fragment **6** is less potent than **5** despite of the addition of the aromatic system. Next, for a positive control, to the sample of **6** an equimolar amount of **BMS-1166** was added and a full recovery of 2D HMQC spectrum of the ^15^N PD-1 was observed.

**Fig. 5.**
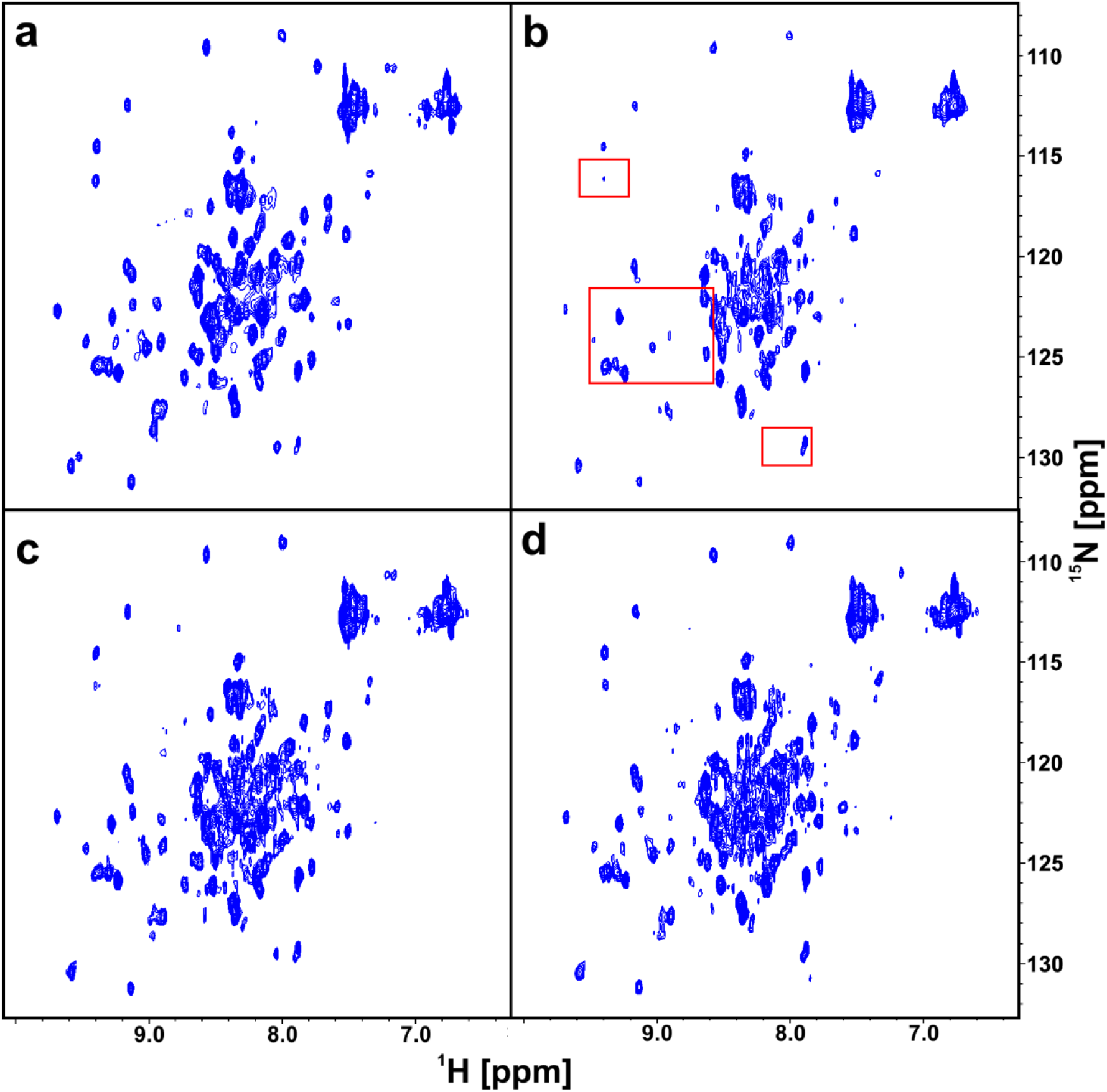
The 2D AIDA NMR for the mutant-PD-1/PD-L1 complexes. The spectra of apo-(N66A)PD-1 (a), the complex of (N66A)PD-1/PD-L1 with marked frames in red the most changing parts of the spectrum (b), the complex of (N66A)PD-1/PD-L1 with **5** (c) and **6** (d) in the molar ratio of the protein complex to the compound 1:1, respectively.

Similar results can also be obtained by analyzing the 1D ^1^H NMR spectra (Supplementary Fig. 18). As a result of the complex formation, significant signal broadening of the spectral lines can be observed. This is clearly visible for the NMR signals of PD-1 with the chemical shifts at −0.2 and −0.7 ppm. After addition of the tested compounds, those signals have partially recovered and significant sharpening of signals between 0.0 and −0.1 ppm were observed. The full recovery of the PD-1 signals was observed only after addition of **BMS-1166**.

We could estimate that the dissociation constant of PD-L1/fragment **5** interaction, which is in the range 50 ± 20 μM. In the case of fragment **6**, for which the w-AIDA-NMR indicated less recovery of the NMR signals, we determine that the K_D_ is around 120 ± 40 μM (the intensities of the recovered resonances are used to obtain an approximate dissociation constant. Errors are quantified from signal-to-noise in NMR spectra). These results correlate with the Homogeneous Time-Resolved Fluorescence (HTRF) assay, where the IC_50_’s for **5** and **6** were determined at 34 μM and above 100 μM, respectively (Supplementary Fig. 19).

### AIDA-NMR on the complex between wt-PD-1 and wt-PD-L1-Long

The K_D_ of the interaction of PD-1 with the construct of PD-L1 that encompasses amino acids 18-239 (PD-L1-Long, 24.3 kDa, is 51 μM) (Supplementary Fig. 11). The wt-PD-1/wt-PD-L1-Long interaction constitutes an example of detecting weak double digit μM to mM binders with the AIDA-NMR on the natively occurring protein-protein interactions.

For the wt-PD-1/wt-PD-L1-Long, we used the smallest fragment of **BMS-1166**, fragment **5**, and tested its dissociating capability for this particular complex (Supplementary Fig. 20). Fragment **5** dissociated the proteins complex, as suggested by the K_D_ of the PPI, and in full agreement with the corresponding data for the w-AIDA-NMR of the mutant N66A of the shorter construct of PD-L1. The PD-1/PD-L1 complex of the latter is more stable in NMR conditions and therefore preferable in the AIDA-NMR experimental settings.

## DISCUSSION

Fragment screening is frequently the first step for the identification and development of molecules that modulate the activity of therapeutic targets. Numerous biophysical methods exist for the identification of fragment hits^60–63^. Among them, the NMR experiment is recognized as a highly robust technique for fragment screening against protein targets^6,8,64–66^. Here, we have described a competition NMR experiment-w-AIDA-NMR - that is sensitive to weak μM-mM interactions and directly shows whether an antagonist releases proteins from their PPI interaction. We believe that w-AIDA-NMR is a valuable complement to the renowned binary ligand-protein SAR-by-NMR assay^6,64,65^, but also to the saturation transfer difference (STD) NMR experiment. For the STD, AIDA-NMR avoids weak points of that experiment as the AIDA-NMR offers checking for compound aggregation and protein instability, two situations leading to false positives. Moreover, by introducing the mutations into non-binding partner, w-AIDA-NMR is performed under physiological conditions as target protein/ligand interface is not compromised. This was further validated using the native PD-1/PD-L1-Long complex that was dissociated by compounds **5** and **6** in a similar manner as the (N66A) PD-1/PD-L1, therefore indicating that the point mutation at the interface of the small molecule non-binding partner indeed does not affect a subsequent target protein/ligand complex. Our NMR results presented here confirmed unambiguously that small-molecule compounds that are able to dissociate the complex between mutant-p53/Mdm2, mutant-p53/MdmX and the mutated PD-1 and PD-L1 could be fished out, and we could estimate the dissociation constant of the targeted protein/fragment interaction. For the PD-1/PD-L1 small-molecule inhibitor **BMS-1166**, we could determine that the minimal fragment of **BMS-1166** responsible for the PD-L1 binding is fragment **5**.

## METHODS

### Purification of Mdm2

The recombinant human Mdm2 (1-125 aa) were cloned into pET11a vector (Novagen) and expressed in *E. coli* BL21 (DE3) RIL (Invitrogen). Cells were grown at 37°C, induced at OD_600_ of 0.6 with 1 mM IPTG and left for expression for additional 4 h at 37°C. Cells were harvested by the centrifugation and the protein was purified from inclusion bodies according to previously described methods^13^. The pellet was resuspended in PBS buffer and sonicated using a macrotip (output control 8, 80%) 5 times for 2 min. Lysate was then clarified by the centrifugation at 60 000 x g for 30 min at 4°C. Inclusion bodies were washed twice with PBS supplied with 0.05% TritonX-100 and centrifuged at 12 000 x g for 25 min. they were later solubilized overnight in 6 M guanidinium hydrochloride, 100 mM Tris-HCl, pH 8.0, with the addition of 1 mM EDTA and 10 mM β-mercaptoethanol at 37°C. Solubilized inclusion bodies were next centrifuged at 60 000 x g for 45 min at 4°C and the resulting supernatant was dialyzed against 4 M guanidinium hydrochloride, pH 3.5, 10 mM β-mercaptoethanol for 6 h. Refolding was performed in 10 mM Tris-HCl, pH 7.0, including 1 mM EDTA, 10 mM β-mercaptoethanol at 4°C then the mixture was supplemented with 1.5 M (NH_4_)_2_SO_4_. Mixture was centrifuged (3000 x g for 35 min at 4°C) before loading on Butylsepharose 6 Fast Flow column. The protein solution was eluted with 100 mM Tris-HCl, pH 7.2, 10mM β-mercaptoethanol. The fractions containing protein were further using SEC column HiLoad 16/60 Superdex 75 prep grade (GE Healthcare) previously equilibrated with 50 mM KH_2_PO_4_, 50 mM Na_2_HPO_4,_ pH 7.4, 150 mM NaCl, 5mM β-mercaptoethanol.

### Purification in native conditions of MdmX, wt-p53, and p53

The recombinant MdmX (18-111 aa) was cloned into pET46Ek/LIC vector (Novagen) and expressed in *E.coli* BL21 (DE3) (Invitrogen). The wt-p53 (1-321 aa) as well as its mutants were cloned into the pET23 vector (Novagen) and expressed in *E. coli* BL21 (DE3) RIL (Invitrogen). Cells were grown in LB medium (100 μg/ml ampicillin) at 37°C and induced with 1 mM IPTG (OD_600_ around 0.6-0.8). The protein expression was performed for additional 12 h at 20°C. Cells were harvested by the centrifugation. The cell pellet was then resuspended in a lysis buffer (50 mM Na_2_HPO_4_, 300 mM NaCl, 10 mM imidazole pH 8.0) and sonicated as described before. After centrifugation at 60 000 x g for 30 min at 4°C, the supernatant was loaded on the Ni-NTA column and incubated for 2 h at 4°C. Loaded protein was washed with buffer: 50 mM Na_2_HPO_4_, 300 mM NaCl, 10 mM imidazole pH 8.0 to elute any unspecifically bound proteins. The taget protein was eluted with buffer: 50 mM NaH_2_PO_4_, 300 mM NaCl, 10 mM imidazole, pH 8.0, relevant fractions were pooled, concentrated and purified by gel filtration in PBS.

### Purification of human wt-PD-1, PD-1 (N66A, Y68A, E135A and N66AY68A), PD-L1 and PD-L1-Long

The human wt-PD-1 (33 – 150 aa, C93S mutation) and mutants (N66A, Y68A, E135A and N66AY68A) were cloned into pET24d (Novagen) while recombinant PD-L1 (18 – 134 aa) and PD-L1-Long (19 – 238 aa) were cloned into pET-21b and pET-28a (Novagen), respectively. Each proteins were expressed in *E. coli* BL21 (DE3) (New England Biolabs). Bacterial cultures were grown overnight at 37 °C in LB or M9 minimal medium and induced at OD600 of ca. 0.8 with 1 mM IPTG. Following inclusion bodies purification was performed using a previously described protocol^59^. Solubilized proteins were refolded with done by a drop-wise dilution into a solution containing 0.1M Tris pH 8.0, 0.4 M L-Arginine hydrochloride, 2 mM EDTA, 5 mM cystamine and 0.5 mM cysteamine (wt-PD-1 and PD-1 mutants) or 0.1 M Tris pH 8.0, 1 M L-Arginine hydrochloride, 0.25 mM oxidized glutathione and 0.25 mM reduced glutathione (PD-L1 and PD-L1-Long). Refolded proteins were next dialyzed 3x against a buffer: containing 10 mM Tris pH 8.0 and 20 mM NaCl and purified using a SEC HiLoad 26/600 Superdex 75 column (GE Healthcare, Chicago, IL, USA) in 25 mM sodium phosphate pH 6.4 with 100 mM NaCl for wt-PD-1 and PD-1 mutants or in PBS pH 7.4 for PD-L1 and PD-L1-Long.

### ITC Measurements

All ITC experiments were performed on a Microcal iTC200 calorimeter according to manufacturer’s protocols. To compensate for the heat generated due to protein dilution a separate experiment was performed, in which a protein solution was injected into a sample chamber with a corresponding buffer. Resulting heat was then subtracted from the final signal as a background.

Data were fitted using Chi^2^ minimization for a model assuming a single set of sites to calculate the binding affinity KD using ORIGIN 7.0. p53-wt and mutants with Mdm2 and MdmX titrations settings were as follow: 20 injections, 25°C cell temperature, 10 μcal/s reference power, 600 s initial delay, 0.012-0.030 mM p53 cell concentration, 0.1-0.3 mM of titrant (Mdm2 or MdmX) syringe concentration, and a stirring speed of 800 rpm. The first injection volume was 0.4 μl with an injection time of 0.2 s, while the rest were 2 μl (4 s injection times). Gaps between injections were set at 150 s and data points were recorded every 5 s. All experiments were performed in duplicate.

### Syntheses

**Nutlin-3a**, compounds **1**, **2**, **3**, and **4** were purchased. Compounds **5** and **6** were synthesized according to the methods described by Guzik et al.^57^.

### Mutagenesis

Site directed mutagenesis of p53 and PD-1 were performed using PCR. The mutagenic primers for p53 (Supplementary Table 1) were designed for QuickChange Site-directed mutagenesis kit (Stratagen). Vectors pET28a with human p53 were used as templates. The mutagenic primers used for PD-1 (Supplementary Table 1) were designed using an inversed PCR approach. Vectors pET24d and pET21b with human PD-1, respectively, were used in the same manner.

### NMR Measurements

Proteins were uniform ^15^N labeled via expression in minimal medium with ^15^NH_4_Cl as the sole nitrogen source. 10% (v/v) of D_2_O was added to the samples to provide lock signal. Spectra were recorded using a Bruker Avance III 600 MHz spectrometer at 300 K equipped with the nitrogen cryo-probe head. By analyzing the line width (relaxation time) of well separated NMR signals we approximated molecular weights of protein populations present in the sample^59^. During the experiment the ^1^H-^15^N signals were monitored by the SOFAST HMQC (Selective Optimized Flip-Angle Short-Transient Heteronuclear Multiple Quantum Coherence)^67^.

### MST assay

The binding affinities between PD-1 (or mutant) with PD-L1 were analyzed using the MST technique. PD-1 and its mutant were fluorescently labelled according the standard labeling protocol of the NanoTemper Protein Labeling Kit RED-NHS (L001-NanoTemper Technologies, Munich, Germany). In MST experiments, the concentration of the labeled PD-1 and mutant was kept constant (20 nM), while the unlabeled PD-L1 were serially diluted in MST buffer (50 mM Tris-HCl buffer, pH 7.6, 150 mM NaCl, 10 mM MgCl_2_ and 0.05% Tween-20) from 416 nM to 183 μM. Samples were premixed and incubated for 2h at RT in dark before loading into capillaries. Data processing is described in the legend of Supplementary Fig. 11.

### Homogenous Time-Resolved Fluorescence (HTRF)

The HTRF assay was performed using the certified Cis-Bio assay kit at 20 μL final volume using their standard protocol as described by Musielak et al.^54^. Measurements were performed on individual dilution series to determine the half maximal inhibitory concentration (IC_50_) of tested compounds. After mixing all components according to the Cis-Bio protocol, the plate was incubated for 2 h at RT. TR-FRET measurement was performed on the Tecan Spark 20M. Collected data was background subtracted on the negative control, normalized on the positive control, averaged and fitted with normalized Hill’s equation to determine the IC_50_ value using Mathematica 12.

### Fluorescence Polarization Assay (FP)

FP competition assay is based on the displacement of p53 mutant peptide called P2 from the complex with Mdm2/X as previously described^68^. All measurements were conducted on Tecan Infinite® 200 PRO plate reader. Buffer formulation was as follow: 50 mM NaCl, 10 mM Tris pH 8.0, 1 mM EDTA, 5% DMSO. FP was determined at λ = 485 nm excitation and λ = 535 nm emission 15 min after mixing all assay components. All tests were performed using Corning black 96-well NBS assay plates at room temperature.

## ACKNOWELEDGMENTS

This research has been supported by the Grants Symphony UMO-2014/12/W/NZ1/00457 and Maestro 2017/26/A/ST5/00572 (to T.A.H.) from the National Science Centre, Poland. We would like to thank Dr. Katarzyna Guzik for excellent support in organic syntheses.

## AUTHOR CONTRIBUTIONS

T. A. H., W. J., and B. M. conceived the study and designed experiments. B. M., W. J., I. R., D. S., K. M.-M., and J. P. carried out the experimental work and data analysis. B. M. and T. A. H. wrote the manuscript. All authors contributed to discussions and revisions of the manuscript.

## ADDITIONAL INFORMATION

Supplementary information

