## Supplementary Materials for "Competition NMR for Detection of Hit/Lead Inhibitors of Protein-Protein Interactions"

**Supplementary Table 1.** Sequence of primers used for mutagenesis of p53 and PD-1.

| **p53 mutant** | **Primer sequence** |
|---|---|
| L22A | Forward: 5’-GAAACATTTTCAGACGCATGGAAACTACTTCCT-3’ |
| | Reverse: 5’-AGGAAGTAGTTTCCATGCGTCTGAAAATGTTTC-3’ |
| L22I | Forward: 5’-GAAACATTTTCAGACATATGGAAACTACTTCCT-3’ |
| | Reverse: 5’-AGGAAGTAGTTTCCATATGTCTGAAAATGTTTC-3’ |
| L22V | Forward: 5’-AAACATTTTCAGACGTATGGAAACTACTTCCT-3’ |
| | Reverse: 5’-AGGAAGTAGTTTCCATACGTCTGAAAATGTTTC-3’ |
| L26I | Forward: 5’-GACCTATGGAAACTAATACCTGAAACAAC-3’ |
| | Reverse: 5’-GTTGTTTTCAGGTATTAGTTTCCATAGGTC-3’ |
| L26V | Forward: 5’-GACCTATGGAAACTAGTTCCTGAAAACAAC-3’ |
| | Reverse: 5’-GTTGTTTTCAGGAACTAGTTTCCATAGGTC-3’ |
| L22IL26V | Forward: 5’-GACATATGGAAACTAGTTCTTGAAAACAAC-3’ |
| | Reverse: 5’-GTTGTTTTCAAGAACTAGTTTCCATATGTC-3’ |
| L22VL26V | Forward: 5’-GACGTATGGAAACTAGTTCCTGAAAACAAC-3’ |
| | Reverse: 5’-GTTGTTTTCAGGAACTAGTTTCCATACGTC-3’ |
| **PD-1 mutant** | **Primer sequence** |
| N66A | Forward: 5’-CTTCGTGCTAGCGTGGTACCGCATG-3’ |
| | Reverse: 5’- CTCTCCGATGTGTTGGAG-3’ |
| Y68A | Forward: 5’-GCTAAACTGGGCGCGCATGAGCCCC-3’ |
| | Reverse: 5’-ACGAAGCTCTCCGATGTG-3’ |
| E135A | Forward: 5’-GCAGATCAAAGCGAGCCTGCGGGC -3’ |
| | Reverse: 5’-GCCTTGGGGGCCAGGGAG-3’ |
| N66AY68A | Forward: 5’-GGCGCGCATGAGCCCCAGCAAC-3’ |
| | Reverse: 5’-CACGCTAGCACGAAGCTCTCCGATG-3’ |

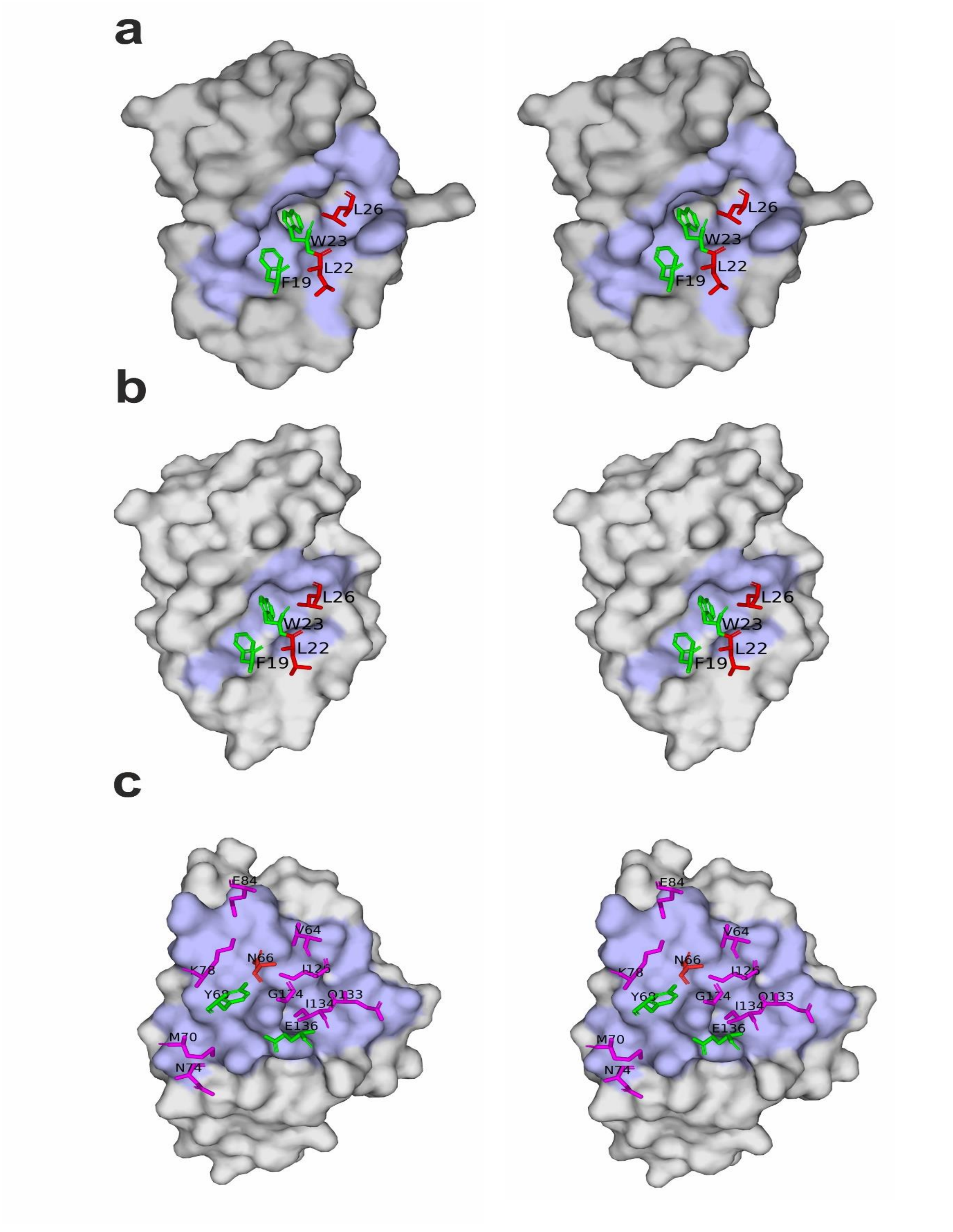


**Supplementary Fig. 1.** Stereo view of a) the Mdm2 complex with p53, b) the MdmX complex with p53 and c) the PD-L1 complex with PD-1. The complexes interfaces are colored blue. Partner amino acid residues that were mutated for this publication are colored red, native crucial residues for the interactions are colored green, while other important residues at the interface are colored purple.

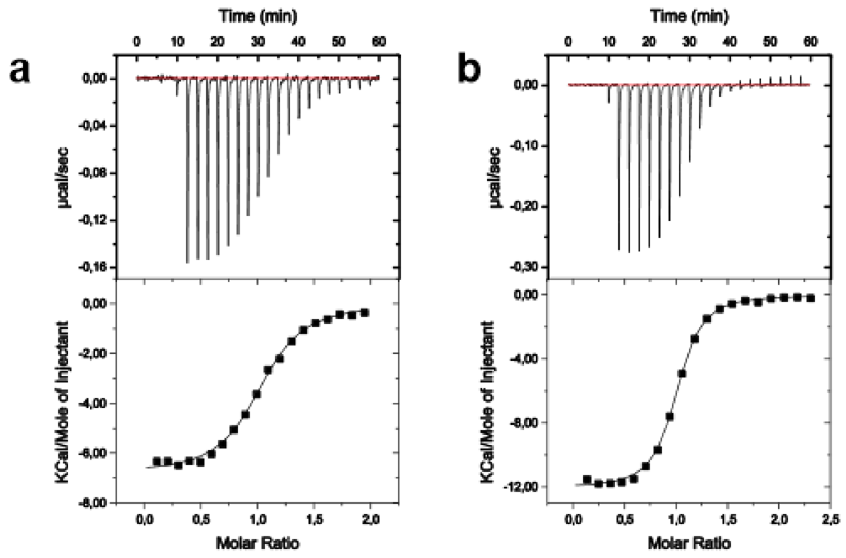


**Supplementary Fig. 2.** Results of the ITC. Binding of p53-wt (1-321) to Mdm2 (a) and MdmX (b).

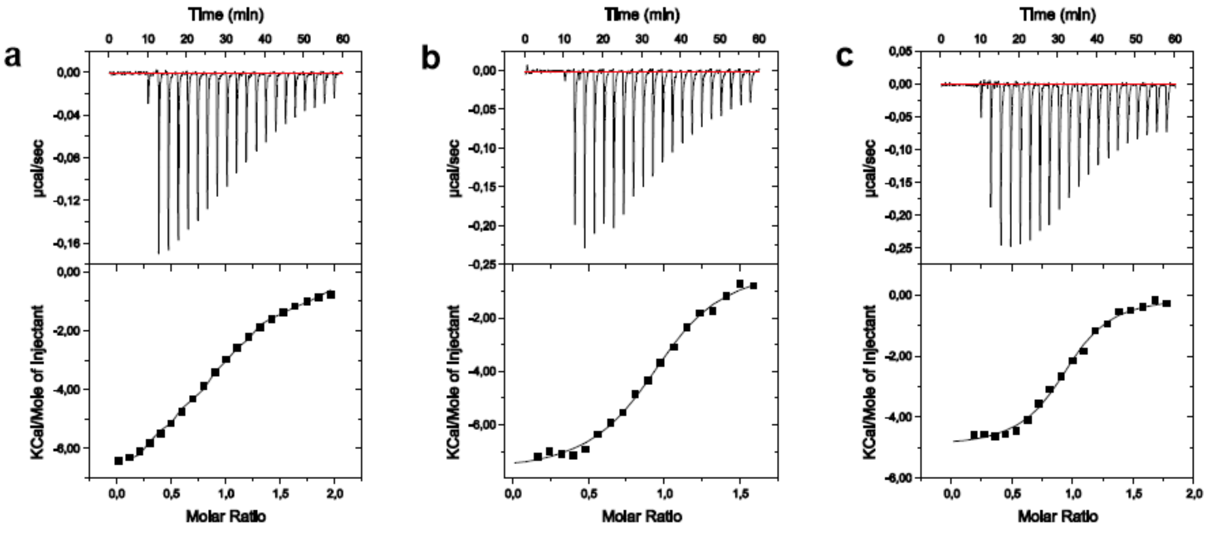


**Supplementary Fig. 3.** Results of the ITC. Binding of p53-wt (1-321) mutants: L22A (a), L22I (b) and L22V (c) to Mdm2.

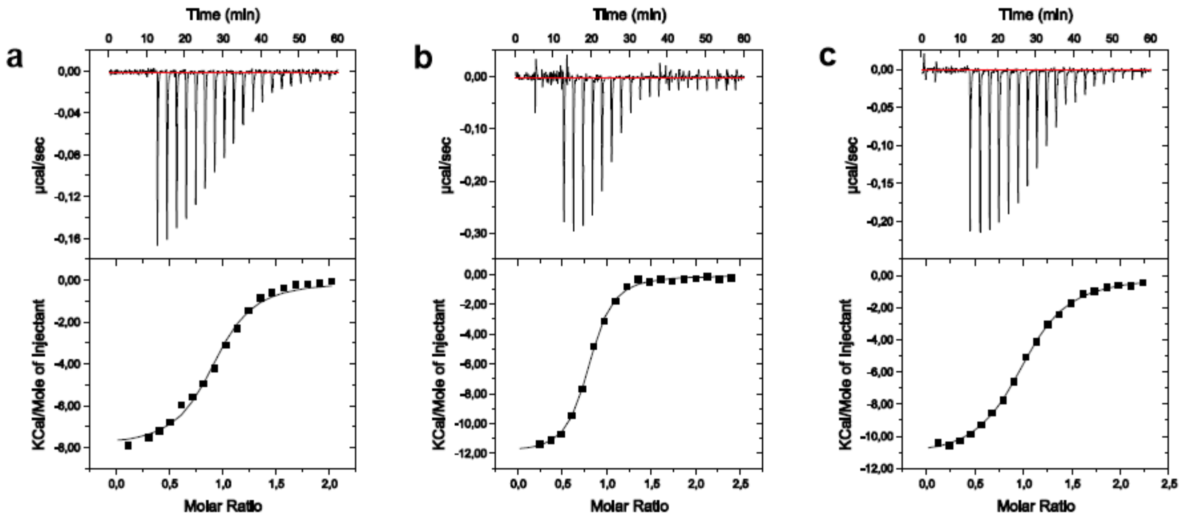


**Supplementary Fig. 4.** Results of the ITC. Binding of p53-wt (1-321) mutants: L22A (a), L22I (b) and L22V (c) to MdmX.

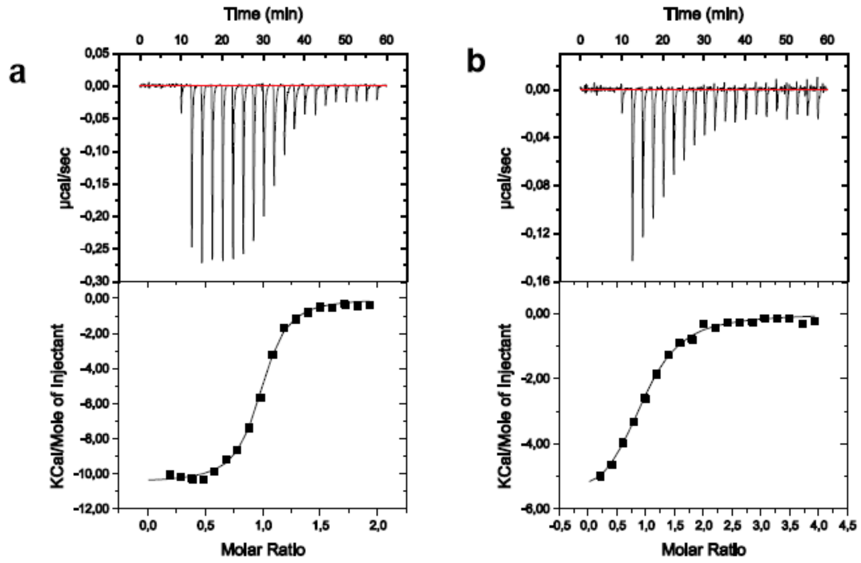


**Supplementary Fig. 5.** Results of the ITC. Binding of p53-wt (1-321) mutants: L26I (a) and L26V (b) to Mdm2.

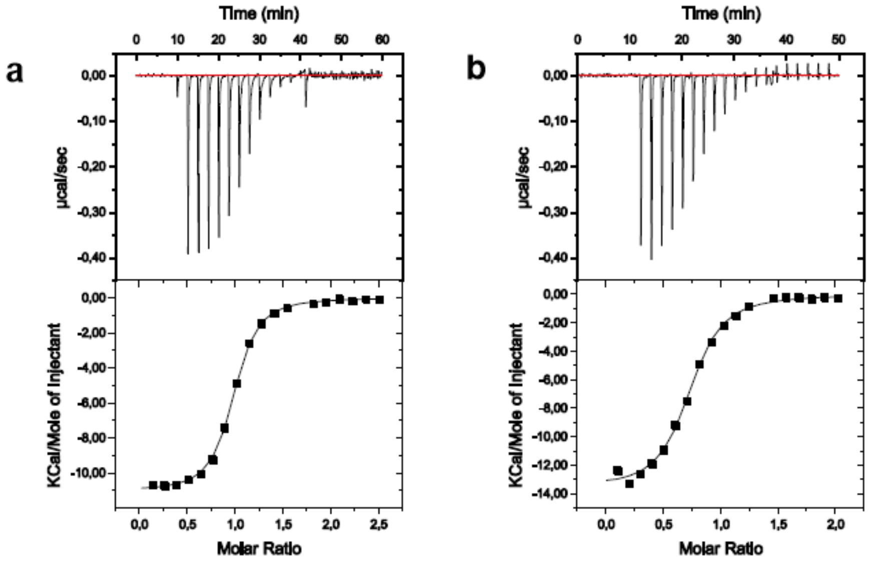


**Supplementary Fig. 6.** Results of the ITC. Binding of p53-wt (1-321) mutants: L26I (a) and L26V (b) to MdmX.

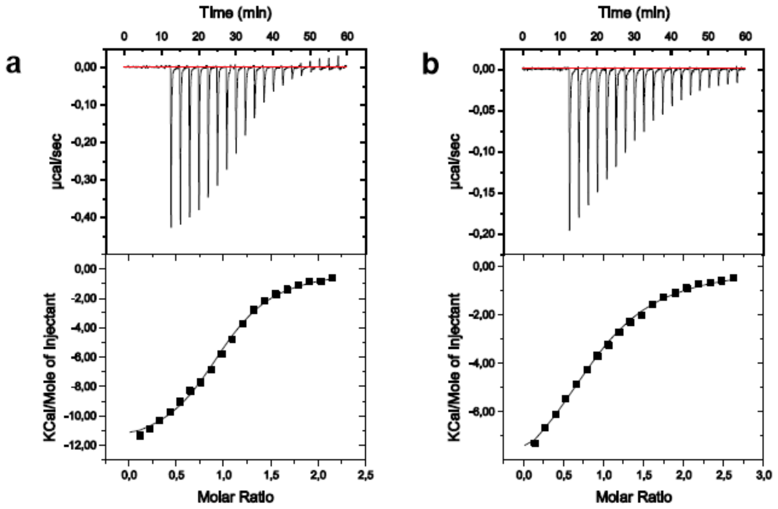


**Supplementary Fig. 7.** Results of the ITC. Binding of different L22VL26V (a) and L22IL26V mutants (b) to Mdm2.

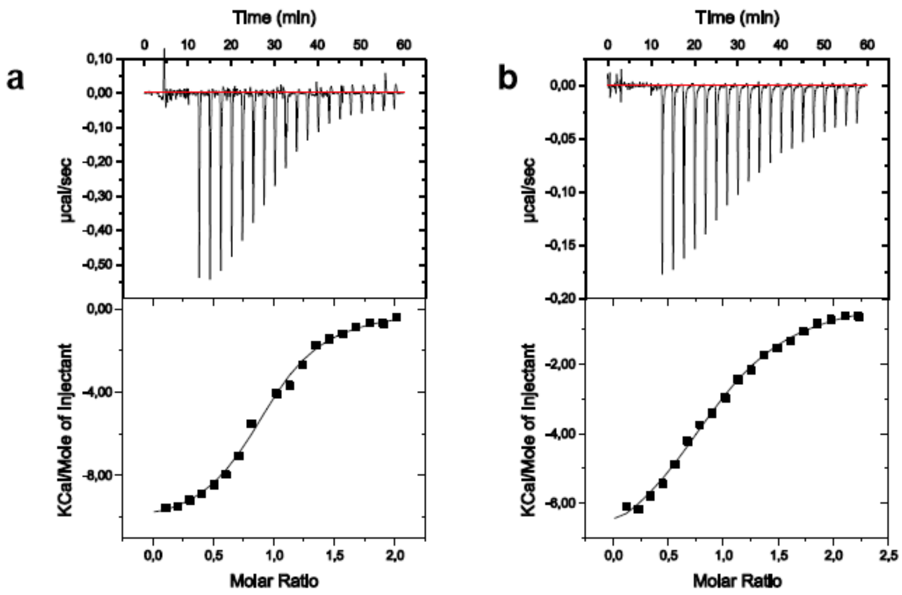


**Supplementary Fig. 8.** Results of the ITC. Binding of different L22VL26V (a) and L22IL26V mutants (b) to MdmX.

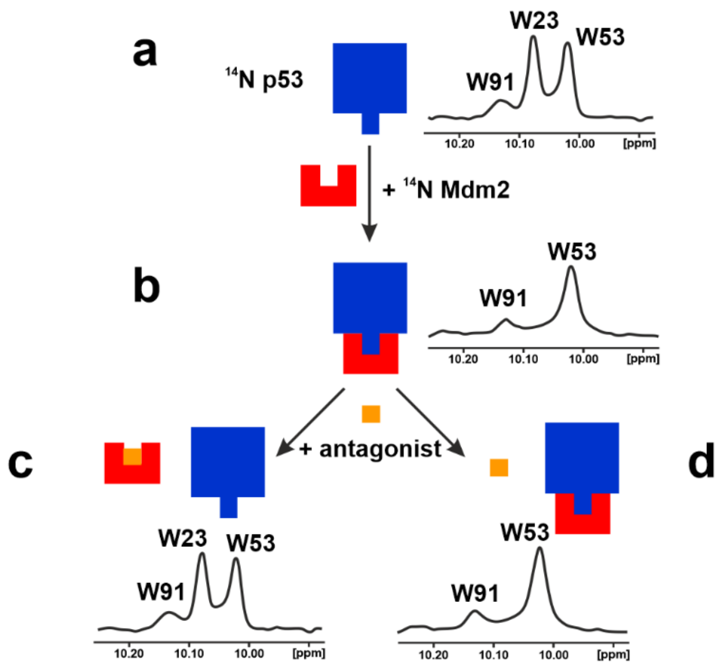


**Supplementary Fig. 9.** A 1D NMR version of the AIDA assay. (a) The 1D proton NMR spectra of the side chain $^{N}H^{\varepsilon}$ protons of tryptophans (W) of free p53 (residues 1-321). The N-terminal domain of p53 contains three tryptophan residues: W23, W53, and W91.[1-2] The side chains of W23 and W53 give rise to sharp lines, because the very N-terminal segment of p53 comprising residues 1-73 has been shown to be very flexible.[3-9] (b) Upon forming the complex with Mdm2 (residues 1-125), the signal of W23 disappears. This is because W23, together with the p53 residues 17 to 26, comprise the primary binding site for Mdm2. Upon binding, these residues participate in a well-defined structure of a large p53-Mdm2 complex, whereas W53 is still not structured when p53 is bound to Mdm2[2,5,6]. Thus, the observed 1/T2 transverse relaxation rate of the bound W23 in the complexes increases thus significantly and broadening of NMR resonances results in the disappearance of this signal in the spectra. (c) Disruption of the Mdm2-p53 interaction results in the release of free p53 and the recovery of the W23p53 NHε signal. The height of W23 peak corresponds to the fraction of free p53 and thus, when total concentrations of the complex and the antagonist are known, the KD of the Mdm2-antagonist interaction can be determined from a single competition experiment.[2] (d) A weak inhibitor does not dissociate the complex.

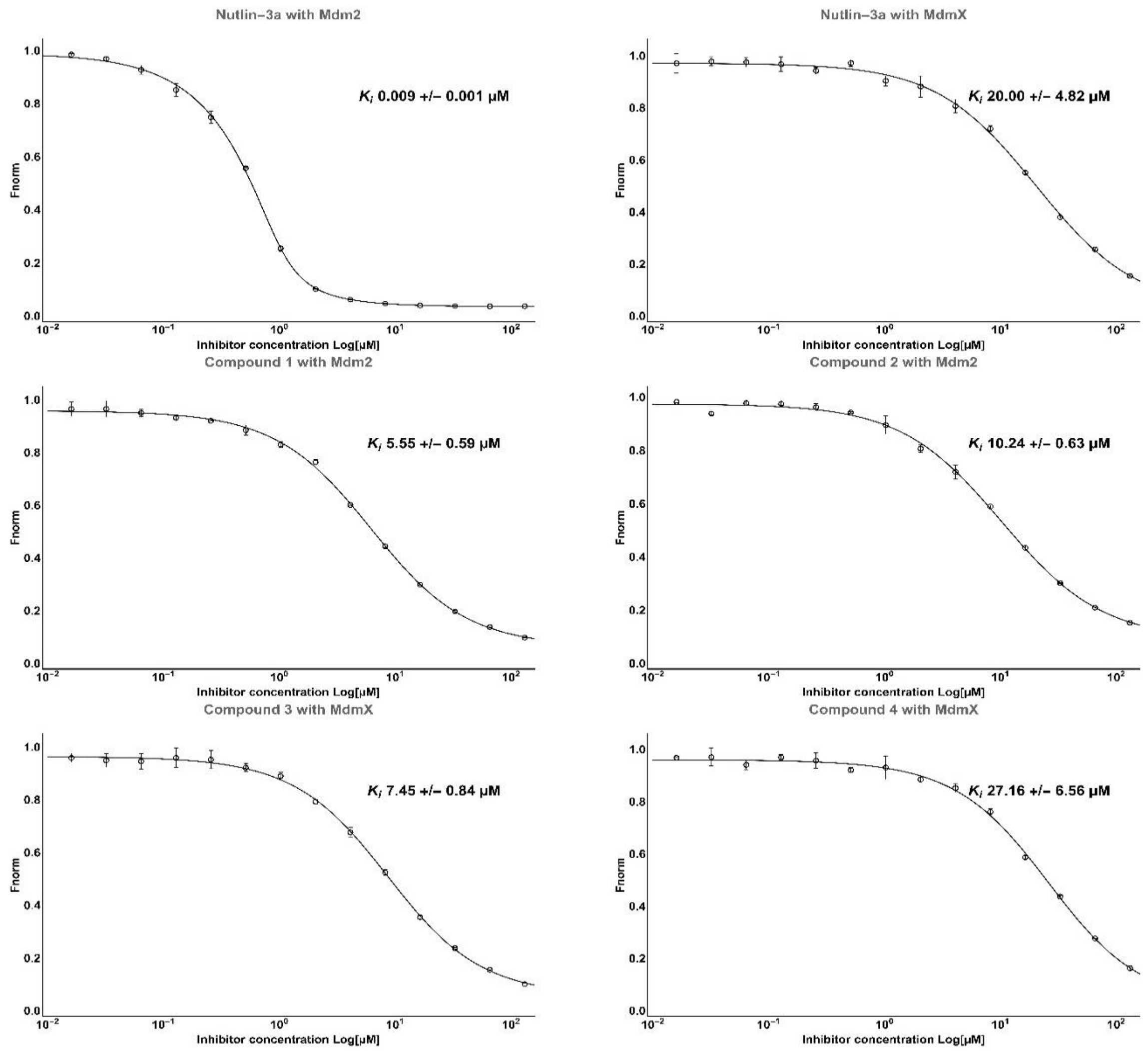


**Supplementary Fig. 10.** Determination of inhibition constants *Ki* from FP assay for Mdm2 protein (middle row) and MdmX (bottom row). Top row is the positive control **Nutlin-3a**. Data is an average of 2 dilution series fitted with the based on the mass balance relationships as described[10].

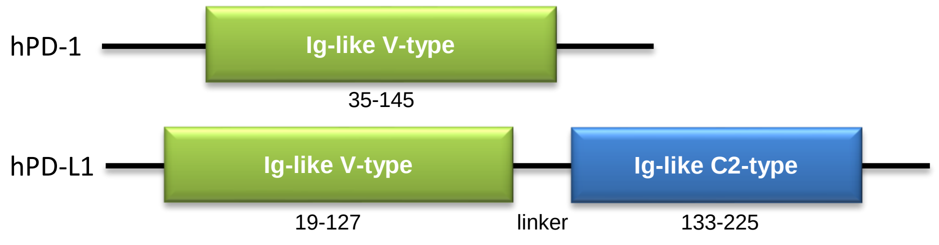


**Supplementary Fig. 11.** Human PD-1 (hPD-1) is a protein of 288 amino acids. The extracellular domain of hPD-1 is 150 amino acid long (amino acids 21-170, 1-20 constitutes the signal peptide). The transmembrane domain is short (amino acids 171-191) and is followed by a cytoplasmic domain of 97 residues (positions 192-288).

Human PD-L1 (hPD-L1) contains 290 amino acids with short both transmembrane and cytoplasmic sequences (residue 239-259 and 260-290, respectively). The ectodomain of hPD-L1 (amino acids 19-238) contains two, the Ig-like V-type and Ig-like C2-type, ca. 100 amino acid domains joined by a short linker. The first N-terminal domain (with the V-type fold) of hPD-L1 is responsible for binding to PD-1. The role of C-terminal domain, characterized by the Ig-like C2-type fold, is unknown (the nomenclature of V- and C2- domain folds relates to the canonical designation of domains within antibodies). It was suggested that it may play a role of a spacer to separate the binding site from the cell membrane. Additionally, crystallographic data showed conformational flexibility between these two domains, which probably allows accommodating to the orientation of PD-1 during the binding[11].

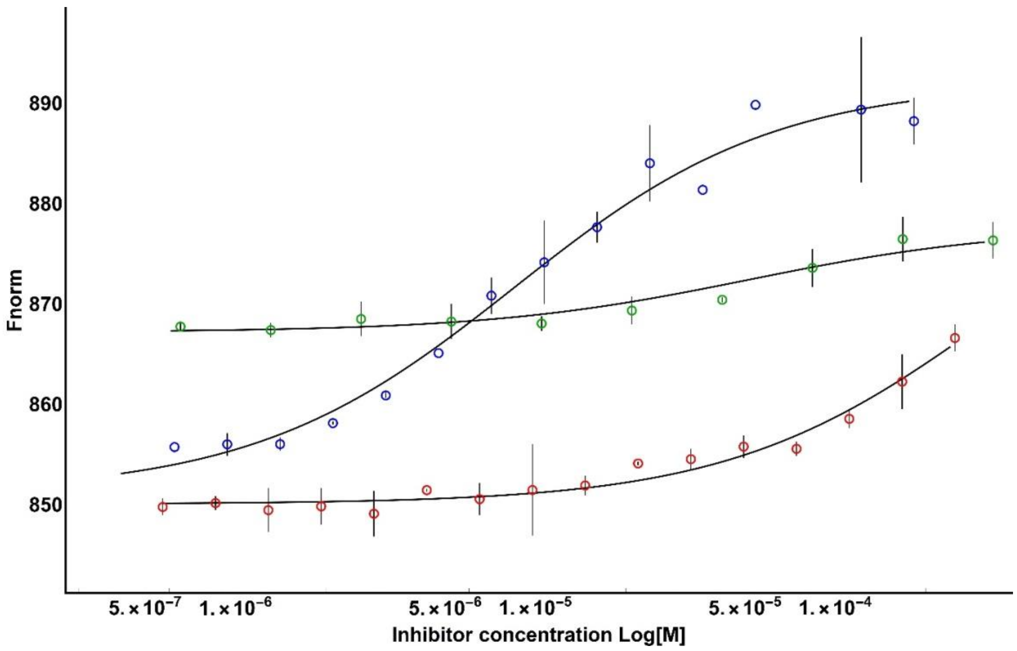


**Supplementary Fig. 12.** Determination of the $K_D$s for PD-1/PD-L1 complex (blue) – 8.0 µM, (N66A)PD-1/PD-L1 complex (red) – over 100 µM and PD-1/PD-L1-Long complex (green) – ca. 51 µM using MST. The MST raw data was fitted with $K_D$ model using dedicated software from Nanotemper and is calculated from the law of mass action:

$$K_D=[A]*[L]/[AL]$$

where [A] is the concentration of free fluorescent molecule, [L] the concentration of free ligand and [AL] is the concentration of the complex of A and L.

The free concentrations of A and L are defined as follow:

$$[A]=[A_0]-[A_L] \text{ and } [L]=[L_0]-[A_L]$$

Where $[A_0]$ is the known concentration of the fluorescent molecule and $[L_0]$ is the known concentration of added ligand.

When combined and rearranged it leads to a quadratic fitting function for [AL]:

$$[AL] = \frac{1}{2} * (([A_0] + [L_0] + K_D) - (([A_0] + [L_0] + K_D)^2 - 4 * [A_0] * [L_0])^{\frac{1}{2}})$$

Since the concentration of fluorescent molecule $[A_0]$ is constant throughout the experiments and the concentration of ligand $[L_0]$ is defined in a dilution series, the signal obtained in the measurement directly corresponds to the fraction of fluorescent molecules that formed the complex $x=[AL]/[A_0]$, which can be fitted with the derived equation to obtain $K_D$.

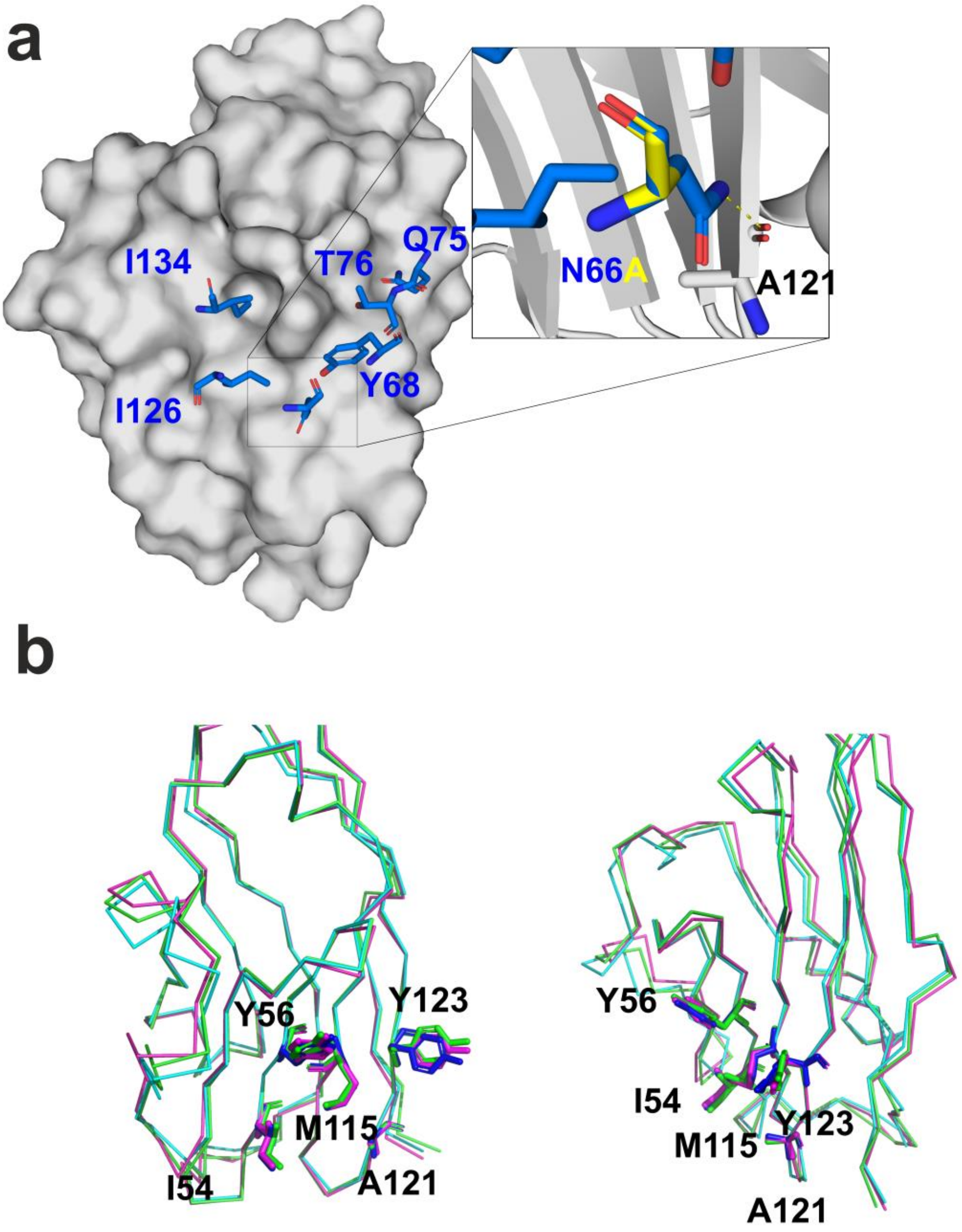


**Supplementary Fig. 13.** The interactions between PD-L1 (gray surface) and selected residues from PD-1 involved in complex formation depicted as blue sticks (PDB ID:4ZQK). (a) The inset presents N66A mutation where the hydrogen bond between the side chain of Asn66 and the carbonyl main-chain oxygen of LAla121is abolished if Asn66 is mutated to alanine (yellow overlay). (b) overlay of the PD-L1 amino acids involved in complex formation from various PDB high resolution crystal structures of PD-1: blue from native complex (PDB ID:4ZQK), green from high affinity PD-1 mutant (5isu) and purple from mPD-1 complex (3SBW).

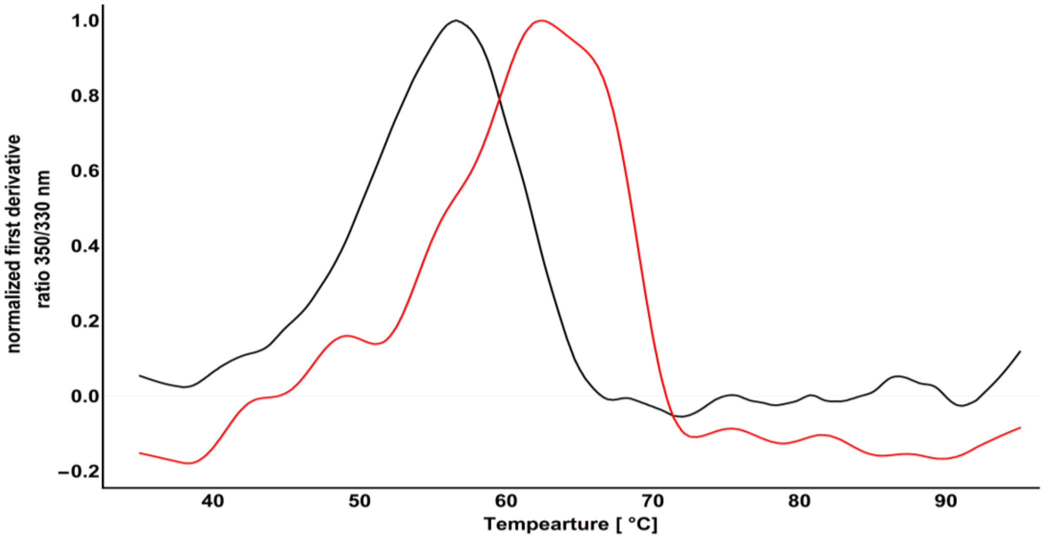


**Supplementary Fig. 14.** The comparison of melting temperatures of wt-PD-1 (black) and (N66A)PD-1 mutant (red). Data represented as a first derivative of the fluorescence ratio at 350/330 nm dependence on the temperature.

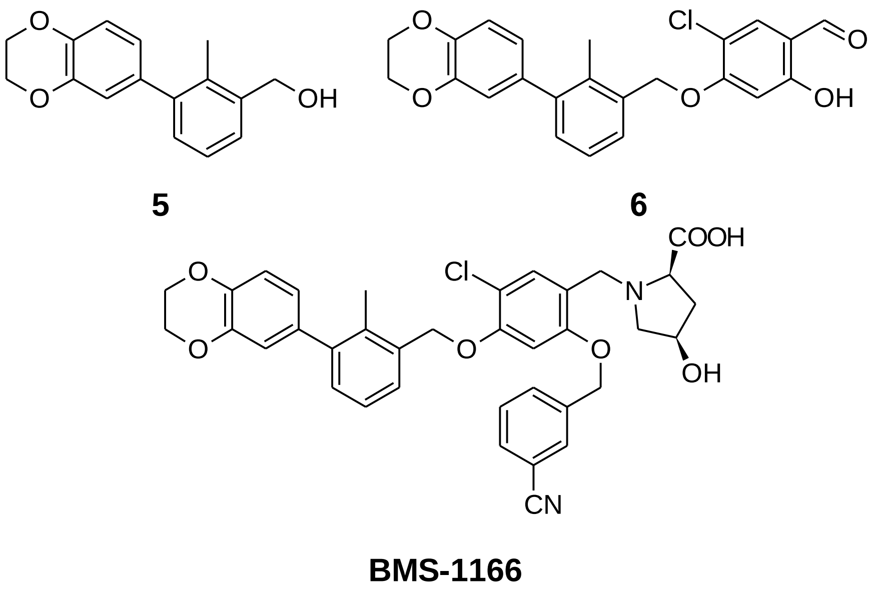


**Supplementary Fig. 15.** The smallest active fragments of **BMS-1166** and **BMS-1166** compound tested for the interaction with PD-L1.

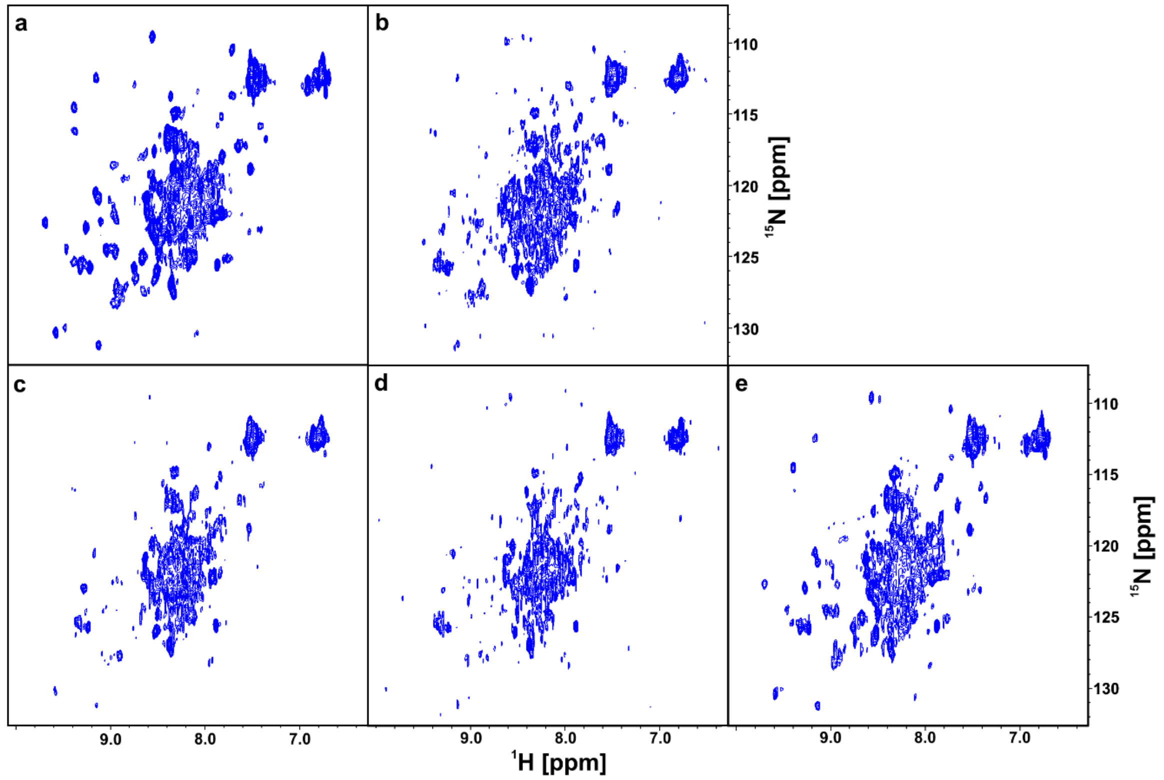


**Supplementary Fig. 16.** $^{1}$H-$^{15}$N HMQC spectra of PD-1 (a), the complex of PD-1/PD-L1 (b), the complex of PD-1/PD-L1 with **5** (c) and **6** (d) in the molar ratio 1:1, and the complex of PD-1/PD-L1 with **5** and **BMS-1166** in the molar ratio 1:1:1 (e).

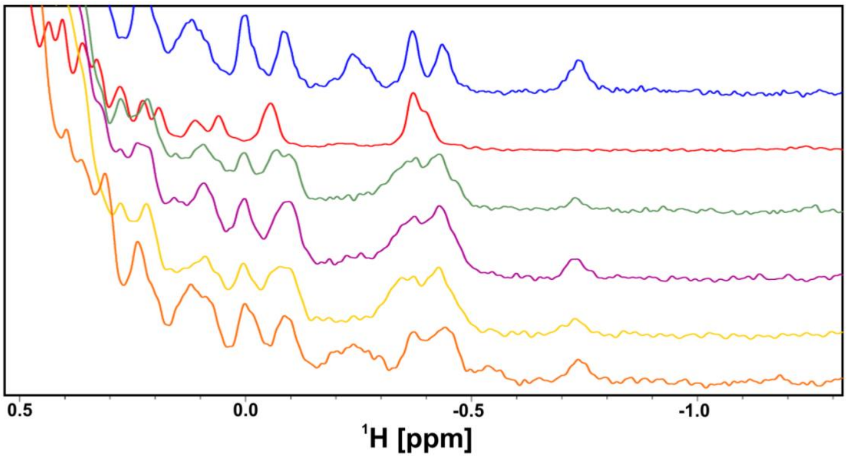


**Supplementary Fig. 17.** The aliphatic part of $^{1}$H NMR spectra of PD-1 (blue), PD-L1 (red), the complex of PD-1/PD-L1 (green), the complex of PD-1/PD-L1 with **5** (purple) and **6** (yellow) in the molar ratio 1:1, the complex of PD-1/PD-L1 with **5** and **BMS-1166** in the molar ratio 1:1:1 (orange).

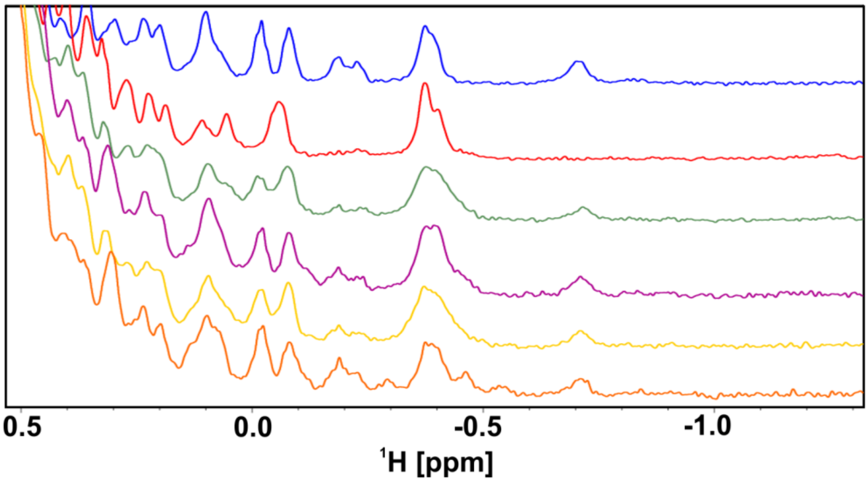


**Supplementary Fig. 18.** The aliphatic part of $^1$H NMR spectra of (N66A)PD-1 (blue), PD-L1 (red), the complex of (N66A)PD-1/PD-L1 (green), the complex of (N66A)PD-1/PD-L1 with **5** (purple) and **6** (yellow) in the molar ratio 1:1, the complex of (N66A)PD-1/PD-L1 with **6** and **BMS-1166** in the molar ratio 1:1:1 (orange).

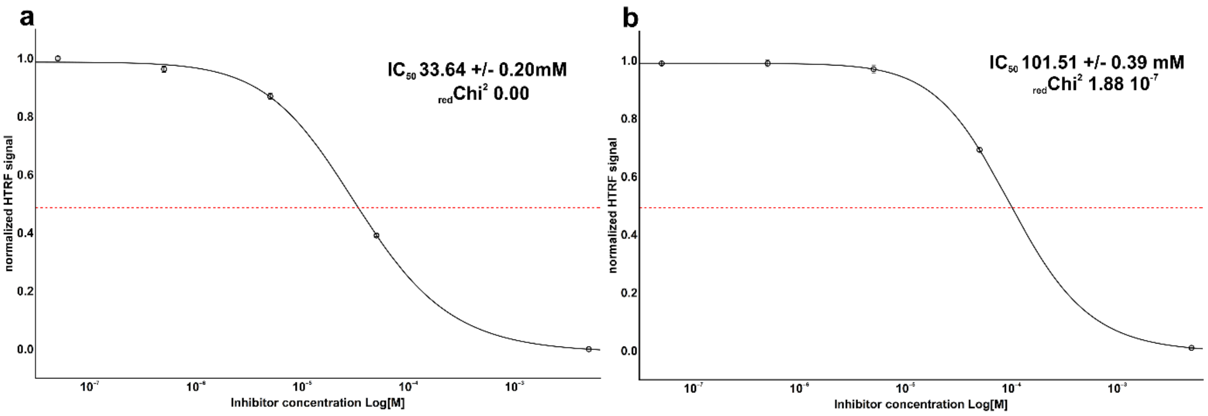


**Supplementary Fig. 19.** The determination of the inhibition constant $IC_{50}$ for compound **5** (a) and compound **6** (b) using HTRF. Raw data was fitted with Hill's model. The goodness of fit is represented in reduced Chi^2.

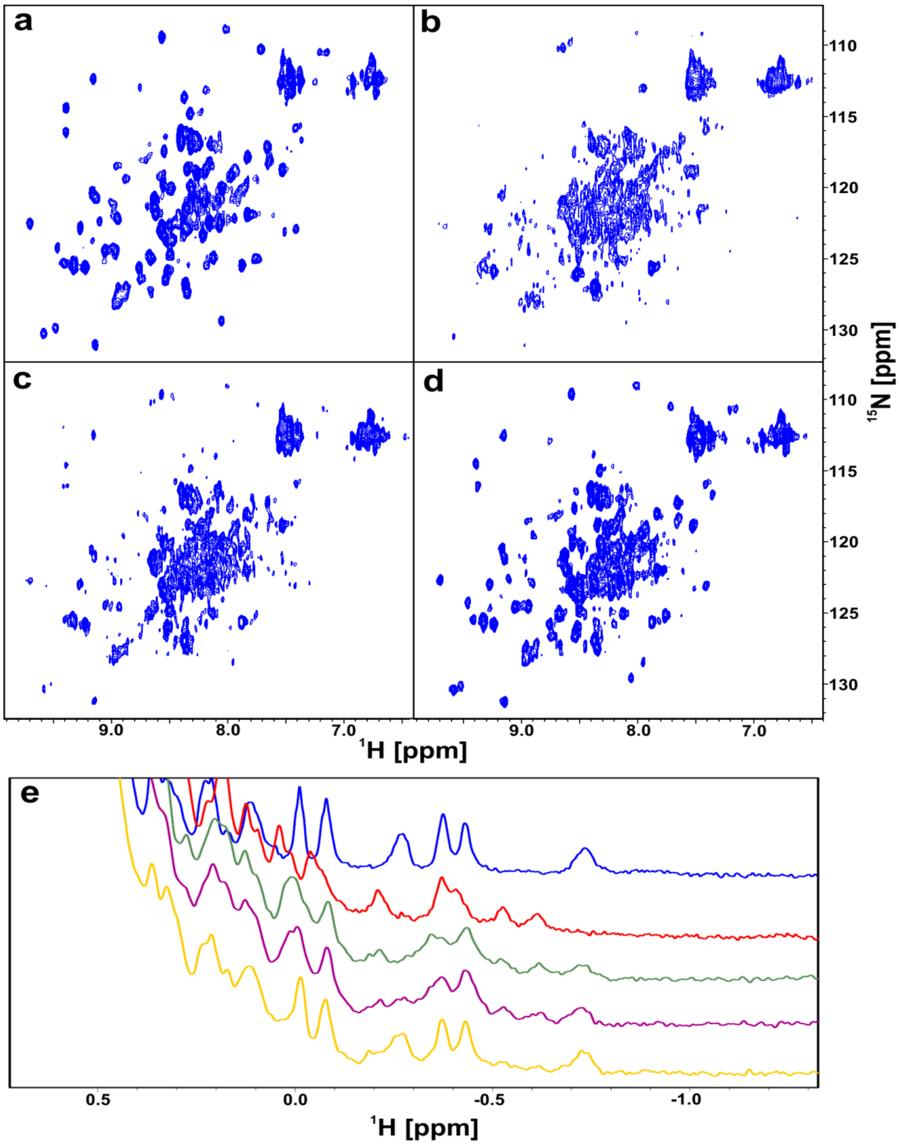


**Supplementary Fig. 20.** $^{1}$H-$^{15}$N HMQC spectra of the $^{15}$N PD-1. (a) Wt-PD-1. (b) The complex of wt-PD-1/wt-PD-L1-Long. (c) The complex of wt-PD-1/wt-PD-L1-Long with **5** in the molar ratio 1:1 of the protein complex and the compound. (d) The complex of wt-PD-1/wt-PD-L1-Long with **5** and **BMS-1166** in the molar ratio 1:1:1 of the protein complex and the compounds. (e) The aliphatic part of $^{1}$H NMR spectra of wt-PD-1 (blue), wt-PD-L1-Long (red), the complex of wt-PD-1/wt-PD-L1-Long (green), the complex of wt-PD-1/wt-PD-L1-Long with **5** (purple) in the molar ratio protein to the compound 1:1, and the complex of wt-PD-1/wt-PD-L1-Long with **5** and **BMS-1166** in the molar ratio 1:1:1 of the protein and the compounds (yellow).
